# Inferring gene-regulatory networks using epigenomic priors

**DOI:** 10.1101/2024.04.23.590858

**Authors:** Thomas E. Bartlett, Melodie Li, Chenyu Song, Yuche Gao, Qiulin Huang

## Abstract

We show improved accuracy in-silico of inference of gene-regulatory network (GRN) structure, resulting from the use of an epigenomic prior network. We demonstrate important use-cases of our proposed methodology by re-analysing 12 datasets from 12 different studies. These include data from cells from human embryos, healthy adult tissue, and cancer, and include single-cell mRNA sequencing data, DNA methylation (DNAme) data, chromatin accessibility data, and histone modification data. We find that DNAme data are very effective for inferring the epigenomic prior network, recapitulating known epigenomic network structure found previously from chromatin accessibility data. Furthermore, we find that inferring the epigenomic prior network from DNAme data reveals candidate TF cis-regulations for at least eight times as many genes, when compared with chromatin accessibility data. When our proposed methodology is applied to real datasets from human embryonic development and from women at risk of breast cancer, we find patterns of differential cis-regulation that are in line with expectations under appropriate biological models, and that may be used to propose hypotheses about pre-cancerous epigenomic changes.

## 1 Introduction

Gene-regulatory network (GRN) models have been shown to be effective at finding specific panels of transcription factors (TFs) that are associated with cell-type identities and with progression of certain diseases (see for example [1–3]). These successful methods of GRN inference are typically based on the target-gene approach [4–6] (Equation 1), which can find important structure in ‘omic networks such as feedback loops [7]. Here, we propose a novel improvement to target-gene GRN inference methodology that builds on our previous work to infer networks from epigenomic data such as DNA methylation (DNAme) data [8, 9], and using advanced regression methods with scRNA-seq (single-cell RNA-seq) and multimodal data [10–12]. Our proposed methodology infers the GRN using advanced regression methods with scRNA-seq (gene-expression) data *following* estimation of an epigenomic prior network. The sequence of these operations is crucial. We show that the accuracy of GRN inference from gene expression data is much improved when using an epigenomic prior network that has been inferred in advance. The epigenomic prior network provides a list of candidate TF regulators that is specific to each target-gene, for subsequent refinement with expression data. This contrasts with alternative strategies, in which the GRN is estimated by combining networks that have been inferred separately and independently from gene expression data and from chromatin accessibility data [1, 2, 5]. Importantly, with our proposed methodology the gene-expression data can originate from a completely independent source to the data that is used to infer the epigenomic prior network. This enables novel analyses specific to tissue and cell-type, by making use of the wealth of genomic and epigenomic data that is now publicly available [13].

A gene-regulatory network (GRN) represents all the gene targets of every TF in a specific cell type. Hence, GRNs are important summaries of the genome-wide gene regulation profile that characterises a particular cellular identity. GRNs have been inferred successfully using methods based on random forests (such as GENIE3 / GRNboost2 [5, 6] as part of the SCENIC / SCENIC+ pipelines [1–3]), as well as alternative approaches based on mutual information [12, 16–19] and recent extensions [20]. However, a genomic network that has been inferred only from gene expression data (such as scRNA-seq data) is more accurately referred to as a gene co-expression network, unless data are included on the interactions between gene-products and DNA [10, 21, 22]. Hence, in order to infer GRNs, data must be included in the inference procedure that allows more precise inference about the interaction of the gene product of the regulating gene (i.e., a TF) with the DNA near the regulated gene (i.e., *cis-regulation*) [1, 3, 11, 12]. Such data often combine the DNA location of cis-regulatory TF binding sites with epigenomic measurements at those locations such as from chromatin accessibility as assessed by ATAC-seq [23]. However, alternative epigenomic data-sources include DNAme data, and ChIP-seq data for histone modifications such as H3K27ac, H3K4me3, and K3K27me3.

In the target-gene approach to GRN inference [4–6], an advanced regression method is used to model the response *x*_*i*_ (the expression-level of gene *i*, Figure 1), as:

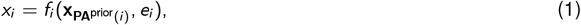

where **PA**^prior^(*i*) are the candidate regulators of gene *i*, 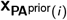 is the vector of expression levels of these candidate regulators, *f*_*i*_ is a (possibly non-linear) function, and *e*_*i*_ is an error or noise term. In our proposed methodology, the epigenomic prior network consists of the candidate regulators of all *p* genes genome-wide, **PA**^prior^(*i*), *i*∈{1, …, *p*}, that are inferred from epigenomic data in combination with TF motif-DNA sequence matching (Figure 1). In our proposed methodology, we first estimate the set of candidate TF regulators **PA**^prior^(*i*) separately for each target-gene *i* from epigenomic data. This is part of an inference procedure where the second step uses advanced regression with gene-expression data to infer the regulators of each *i* from the candidates provided by **PA**^prior^(*i*) (Equation 1, Figure 1). State-of-the-art methods for target-gene GRN inference such as GENIE3 / GRNboost2 [5,6] (as part of the SCENIC / SCENIC+ pipelines [1–3]) do not use candidate sets of regulator TFs **PA**^prior^(*i*) that are specific to target-gene *i*. This makes it more difficult for the advanced regression method to estimate *f*_*i*_ (Equation 1), because the non-target-specific set of candidate TFs must be much larger (of the order of 100s or 1000s) than the target-specific sets (of the order of 10s of TFs). Hence, we use the same type of advanced regression (random-forests) as algorithms such as GENIE3 and GRNboost2 do, but with the novel improvement that in the advanced regression step to estimate *f*_*i*_, we use much smaller sets of candidate TFs **PA**^prior^(*i*) that are specific to each target-gene *i*. These target-specific candidate TF regulators **PA**^prior^(*i*), *i*∈ *{*1, …, *p*}, are encoded in the epigenomic prior network.

**Figure 1.**
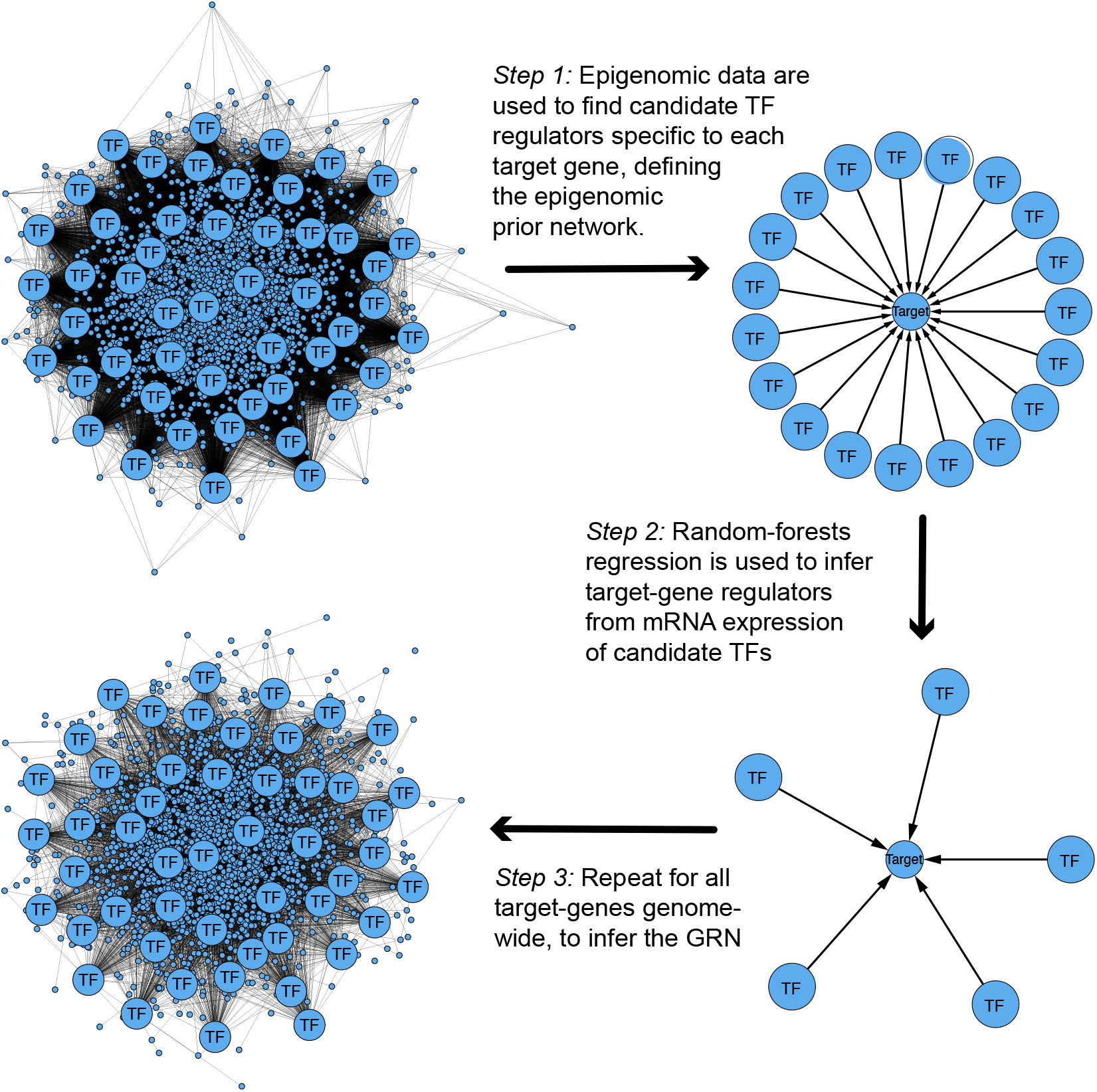
Proposed gene-regulatory network (GRN) inference procedure, for both real data and simulation study. Step 1: Epigenomic data (such as ATAC-seq / LiCAT-seq, H3K27ac, DNAme, data) are used to infer the prior network structure, by analysing epigenomic patterns at cis-regulatory TF binding sites [14, 15]. This provides a set of candidate TF regulators specific to each target-gene, based on whether a candidate gene-regulation is physically realistic. Step 2: An advanced regression method such as random-forests is used to infer the cis-regulators of the target-gene from the specific set of candidate TFs defined in the epigenomic prior network. Step 3: The global gene cis-regulatory network is constructed, by combining the local networks that were inferred around each target-gene [4] in the previous steps.

It is well known that changes in DNAme patterns precede cancer formation [24, 25]: these pre-cancerous changes are sometimes called ‘epigenomic field-defects’ [24, 26] or ‘epigenetic field cancerisation’ [27]. We have previously found that that unique epigenomic patterns distinguish cancer [28], and can be used to predict clinical outcomes [29], and it is well established that DNAme patterns enable different tissues and cell types to be distinguished from one another [30]. Changes in DNAme rates at regulatory DNA regions enable changes in gene regulation [31] and therefore allow changes of phenotype [32], motivating the methodology we propose here to systematically analyse genome-wide DNAme rates at putative binding sites of specific TFs. Recently, it has become clear that a large amount of the observed variation in DNAme patterns in bulk-tissue genomic data can be explained by the variation in the proportion of cells from different lineages that comprise the tissue sample [33]. Hence bulk-tissue DNAme data contain important information about the cell type composition of the tissue, and this finding can also be used as a clinical biomarker, by tracking cells-of-origin of cancer [27, 34]. In the methodology proposed here, we infer GRNs using DNAme data from specific cell types (rather than from a mixture of cells of different lineages), to allow inferences about the functional effects of DNAme changes at cis-regulatory TF binding sites. We use these cell-type specific DNAme data to infer genomic networks, showing how DNAme data can be used to infer the structure of the epigenomic prior network.

It has previously been established that DNAme data are very effective for GRN inference in combination with gene expression data [35]. We have found that an epigenomic prior network (for example based on DNAme data) is essential to achieve the most accurate inference from the advanced regression methods applied to gene expression data such as scRNA-seq data (Figures 1, 2, 3, and 7, and Table 1). We show that the epigenomic prior networks we infer recapitulate well-validated TF cis-regulations of target-genes obtained using the widely used and well validated software packages pycistarget [1, 3, 15] and RGT-HINT [36]. We then infer GRNs from gene-expression (scRNA-seq) data in combination with these epigenomic prior networks, in the contexts of data from human embryonic development, and from studies of women at risk of breast cancer. We find GRNs characteristic of different lineage choices and disease at-risk states, with patterns of differential gene-expression that are in line with expectations under appropriate biological models. However, we note that novel findings from computational GRN inference remain hypotheses until they are experimentally validated. Therefore to give confidence that our proposed methodology leads to verifiable inferences, we use the most significant inferences from our methodology as case studies, showing that they correspond to existing knowledge that is well supported by previous studies.

**Table 1.** Results of the ablation study. Significance-test p-values and t-statistics show that the z-scores calculated in the ablation study are significantly greater than 0 over all target-genes genome-wide, consistently across cell-types and conditions, demonstrating the added value of the epigenomic prior network to the GRN inference.

|  | *t*-statistic | *p*-value |
| --- | --- | --- |
| luminal progenitor WT | 20.7 | <0.001 |
| luminal progenitor BRCAmut | 13.5 | <0.001 |
| luminal mature WT | 21.1 | <0.001 |
| luminal mature BRCAmut | 13.6 | <0.001 |
| basal WT | 19.3 | <0.001 |
| basal BRCAmut | 6.52 | <0.001 |
| fibroblast type1 WT | 21.4 | <0.001 |
| fibroblast type1 BRCAmut | 11.4 | <0.001 |
| fibroblast type2 WT | 34.6 | <0.001 |
| fibroblast type2 BRCAmut | 26.3 | <0.001 |

**Figure 2.**
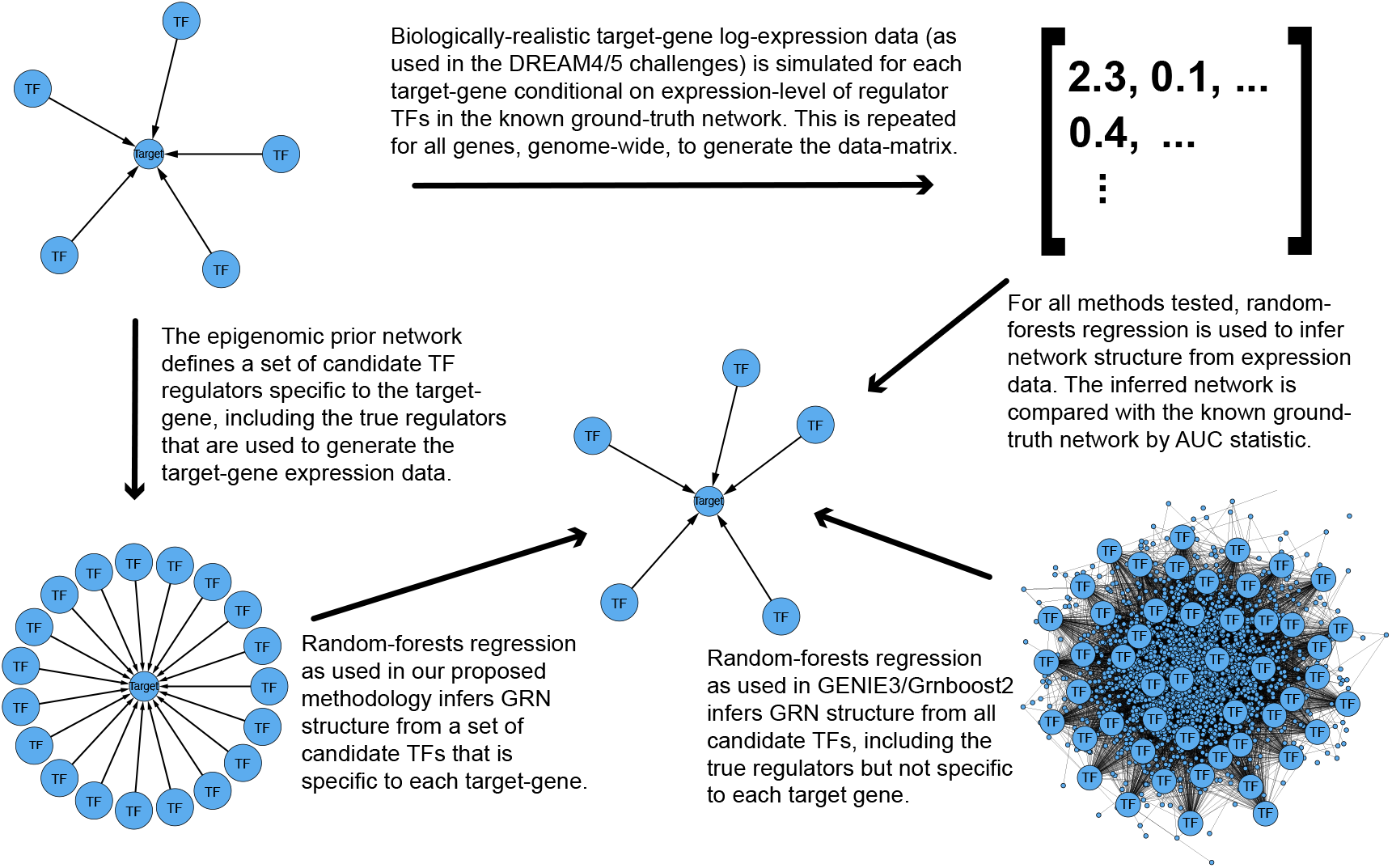
Biologically-realistic gene-expression data (as used in the DREAM4/5 challenge) was generated based on known ground-truth network structure. The prior network provides target-specific lists of candidate TF regulators, which were generated by combining the list of true regulators defined by the ground-truth network, with a random sample of other regulators found in the ground-truth network. Random-forests regression was then applied in two ways. Firstly, to the target-specific lists of candidate regulators defined in the prior network, as in our proposed methodology (Figure 1). Secondly, to the full list of all regulators in the prior network, i.e. not target-specific as required in GENIE3/Grnboost2. The findings are shown in Figure 3

**Figure 3.**
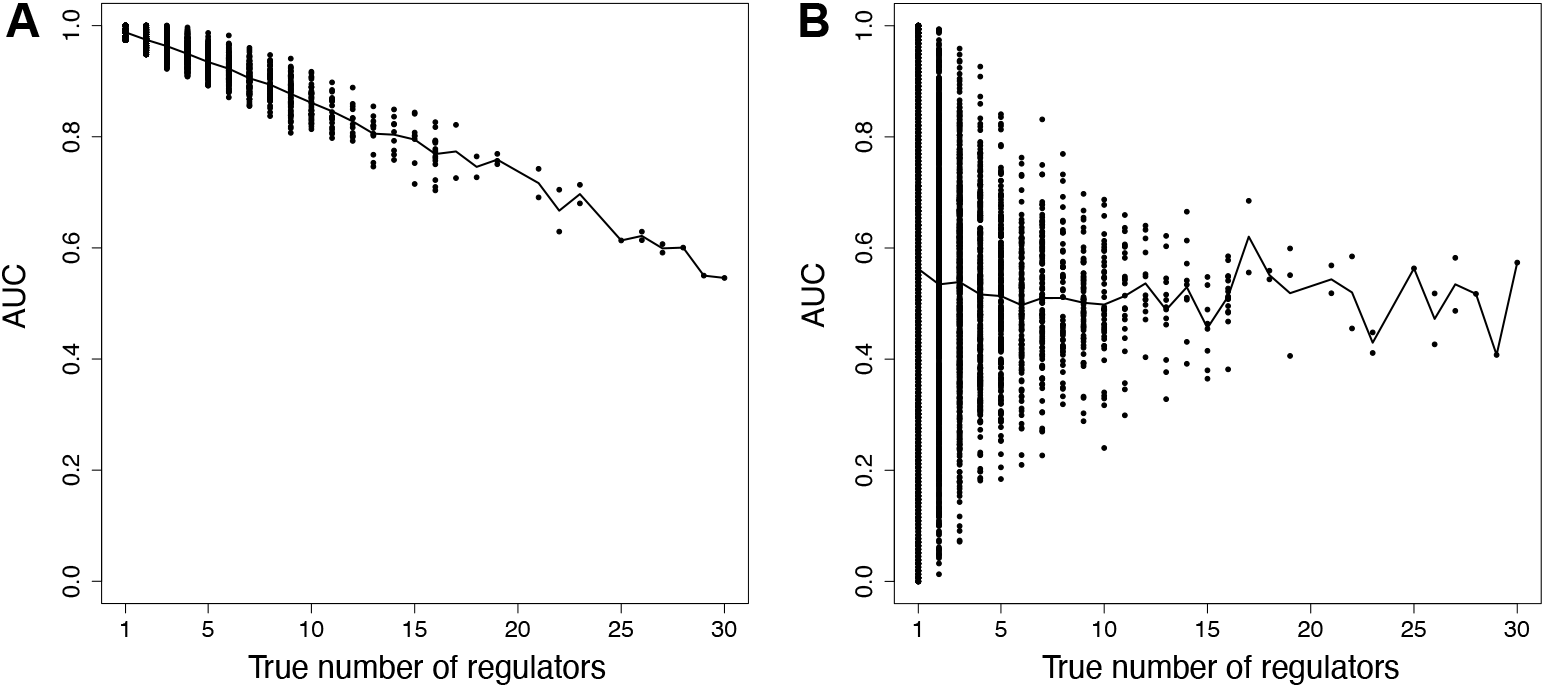
Results from GRN inference using random-forests regression with biologically-realistic gene-expression data generated from known ground-truth network structure. (a) Random-forests target-gene GRN inference implemented as in our proposed methodology (Figure 1). The prior network structure provides a target-specific set of candidate regulators, from which random-forests regression chooses. (b) Random-forests target-gene GRN inference, implemented with GENIE3. Random forests regression infers the target regulators by choosing from all candidate regulators that are found in the prior network, i.e., not from a list that is specific to each target gene.

## 2 Results

### 2.1 Using prior network structure from epigenomic data improves performance of regulatory network inference from gene expression data

We found that the epigenomic prior network allows the advanced regression method (Equation 1) to make more accurate predictions of gene regulations, in the context of biologically-realistic data generated from known network structure (Figure 2). When there are fewer candidate regulators of a target-gene, we found improved accuracy (assessed by AUC) identifying known gene regulations, as shown in Figure 3. These results are based on data generated from known network structure, that was simulated using realistic differential-equations models of chemical reaction kinetics [37] (Section 4). We hence use the AUC statistic to test how well the network edge strengths found by advanced regression, can predict the known ground-truth network structure (that is not otherwise used in the network inference). It can be seen from Figure 3 that there is an increase in accuracy of GRN inference resulting from the reduction in the number of candidate TF regulators that the advanced regression method must choose from, from the order of 100s or 1000s down to the order of 10s. This reduction in number of candidate regulators is enabled by the epigenomic prior network.

For each target-gene, the epigenomic prior network defines a list of candidate TF regulators of the target-gene that are specific to that target-gene. This is based on whether the regulation of target by TF is physically realistic. The target-gene specific list of candidate regulators is typically a few 10s of TFs, making it much easier for the advanced regression method to identify the correct TF regulators of the target-gene (Figure 3a), than it would be from a much longer list of candidate TFs that is not target-specific, as required by GENIE3 / GRNboost2 and similar [1–3, 5, 6]. With existing methods such as GENIE3 / GRNboost2, the full list of TFs (typically of the order of 100s or 1000s of TFs) must be specified as candidate regulators of each target-gene, meaning that it is much more challenging for those methods to find the true TF regulators of the target-gene (Figure 3b). Hence, the epigenomic prior network allows greater accuracy of GRN inference from advanced regression methods such as random forests [38], when used with single-cell gene expression data in a target-gene GRN inference procedure [4].

### 2.2 Recapitulating known epigenomic network structure using DNA methylation data

For the human embryonic development LiCAT-seq (low-input chromatin accessibility) dataset [39], we first verified whether TF-motif cis-regulations that were found using RGT-HINT [36] in previous work [12] would also be discovered from the same LiCAT-seq data by our proposed modified-lever-cistarget (MLC) method for epigenomic prior network inference (Section 4). We found 170 TFs that were present in both analyses, and found that the corresponding regulatory regions overlapped significantly for 168 out of these 170 TFs (Figure 4a). We also applied pycistarget [1, 3, 15] to the same LiCAT-seq human embryonic development dataset [39], finding 124 signficant TF motifs: a large number of these 124 motifs found by pycistarget were also found to be significant according to the MLC method (Figure 4b). Hence, we were able to validate the TF-motif cis-regulatory inferences found by MLC from the LiCAT-seq human embryonic development dataset [39], in comparison with TF-motif cis-regulations that were found from same dataset using RGT-HINT [36] in previous work [12], and also in comparison with inferences from the pycistarget method which is part of the SCENIC+ pipeline [3].

**Figure 4.**
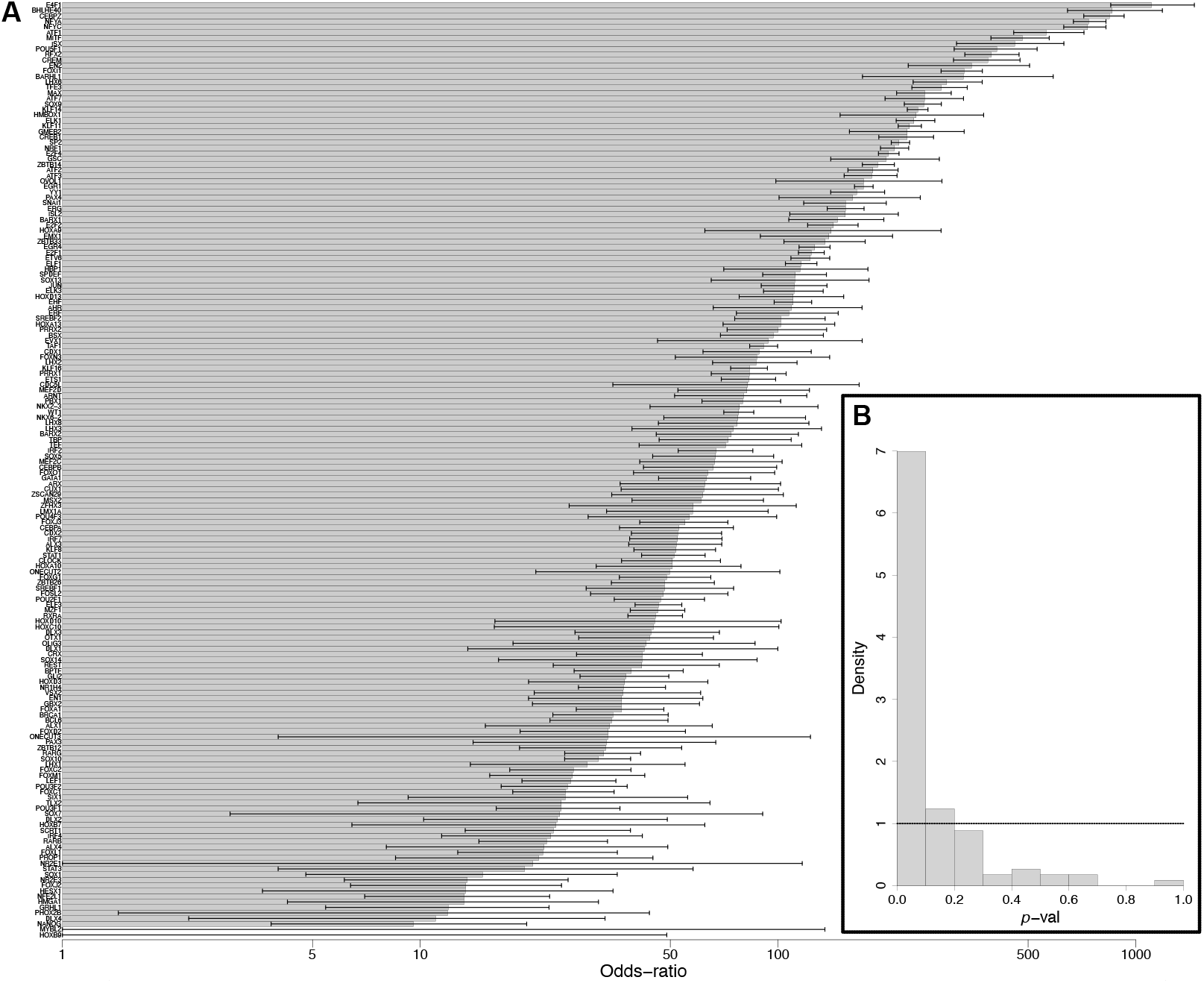
(a) Tests of overlap of the DNA regions inferred as regulated by 170 TFs in the inner cell mass (ICM) according to both the modified-lever-cistarget (MLC) method and the RGT-HINT method [36] (as published previously [12]). Bars show odds-ratios (on a logarithmic scale), with whiskers showing 95% C.I.s. The corresponding significance p-vals (Fisher’s exact test) are provided in Table S1. (b) The 124 motifs inferred when the pycistarget software is applied to the same LiCAT-seq data are are tested for significance according to the MLC method (z-test). The horizontal dashed line at density of 1 indicates the expected density for a uniform distribution of p-values, which would be expected if there were no correspondence between the TF-motif inferences from pycistarget and MLC for this LiCAT-seq dataset. LiCAT-seq is low-input chromatin accessibility data.

Applying the MLC method to the LiCAT-seq human embryonic development dataset [39] allows an epigenomic prior network to be inferred. Equivalently, applying our proposed MethCorTarget methodology (Section 4) to the DNAme human embryonic development dataset [40] allows an epigenomic prior network to be inferred. These epigenomic prior networks inferred from different epigenomic data sources are expected to contain overlapping and similar information. We verified this by testing the overlap between the TF motif-gene cis-regulations inferred by MLC as applied to the LiCAT-seq human embryonic development dataset [39], and the cis-regulations inferred by MethCorTarget applied to the DNAme human embryonic development dataset [40]: we found a highly significant overlap, odds-ratio OR = 13.3 (12.9 − 13.8) (*p* < 0.001, Fisher’s exact test). This large overlap between the lists of cis-regulations inferred from the different data-sources illustrates the interchangeability between the epigenomic prior networks obtained for the same cell type from DNAme data and from chromatin accessibility data. We also note that epigenomic prior network structure inferred from DNAme may be more useful in certain applications, because it provides information about the regulation of more than eight times as many genes (23575 regulated genes found by MethCorTarget from DNAme data, compared to 2650 for MLC from LiCAT-seq data).

### 2.3 Inferring GRNs in epiblast and primitive endoderm cells of the human embryo

Epiblast (Epi) and primitive endoderm (PrE, or hypoblast) cells are the main cell types of the inner cell mass (ICM) of the blastocyst of the human embryo at 5-7dpf (days post fertilisation) [41]. We have previously used ICM chromatin accessibility data to inform inference of GRNs from scRNA-seq data in Epi and PrE [12]. We found that our epigenomic prior network obtained from DNAme data overlaps strongly with that obtained from chromatin accessibility data, but covers a much larger number of target-genes. We therefore proceeded to use the epigenomic prior network obtained from DNAme data as a valid epigenomic prior network to improve GRN inference from the human embryonic development scRNA-seq dataset [42]. We inferred GRNs from scRNA-seq data as described (Figure 1 and Section 4) for both Epi and PrE cell types, using the scRNA-seq dataset cell type definitions shown in Figure S1 and Figure S2, and using the epigenomic prior network obtained from the DNAme human embryonic development dataset (ICM cells) [40].

An example of cis-regulations inferred as part of the GRN for Epi and PrE is shown in Figure 5. The GRN was inferred by filtering candidate cis-regulations found as part of the epigenomic prior network from DNAme data for ICM cells, by variable selection using advanced regression methods and scRNA-seq data for Epi and PrE (according to the procedure outlined in Figure 1). Figure 5 shows DNAme rate profiles for the 103 ICM cells (which include Epi cells and PrE cells) in the DNAme human embryonic development dataset [40] for the *SOX17* gene. The *SOX17* gene is a well known marker gene of PrE cells in contrast with Epi cells [44], and hence we have used it as an example here. In Figure 5, changes in the DNAme profile are visible in regions corresponding to previously-published and validated LiCAT-seq peaks [12, 39], as well as around regulatory regions corresponding to the motifs of candidate TF-regulators of *SOX17*. Figure 5b also shows how in epiblast cells, the epiblast marker gene *NANOG* (Figure S1 and Figure S3) negatively regulates *SOX17* (Figure S2 and Figure S3), which is consistent with *NANOG* maintaining Epi identity by repressing PrE identity [43]. Hence, this example demonstrates how findings from our proposed methodology are in line with expectations under a well-known biological model in human embryonic development. Also interestingly, Figure 5 illustrates how the DNAme rates at these regulatory regions are highly variable, a feature we note is also of interest in understanding cancer formation and cancer predisposition [28, 29]. There are well established similarities between gene-regulatory processes that have been observed in embryonic development and in cancer [45], and we hypothesise that phenotypic plasticity is represented by the variability in methylation rates that is visible in Figure 5 at CpG loci around the motifs of the candidate TF cis-regulators of *SOX17*.

**Figure 5.**
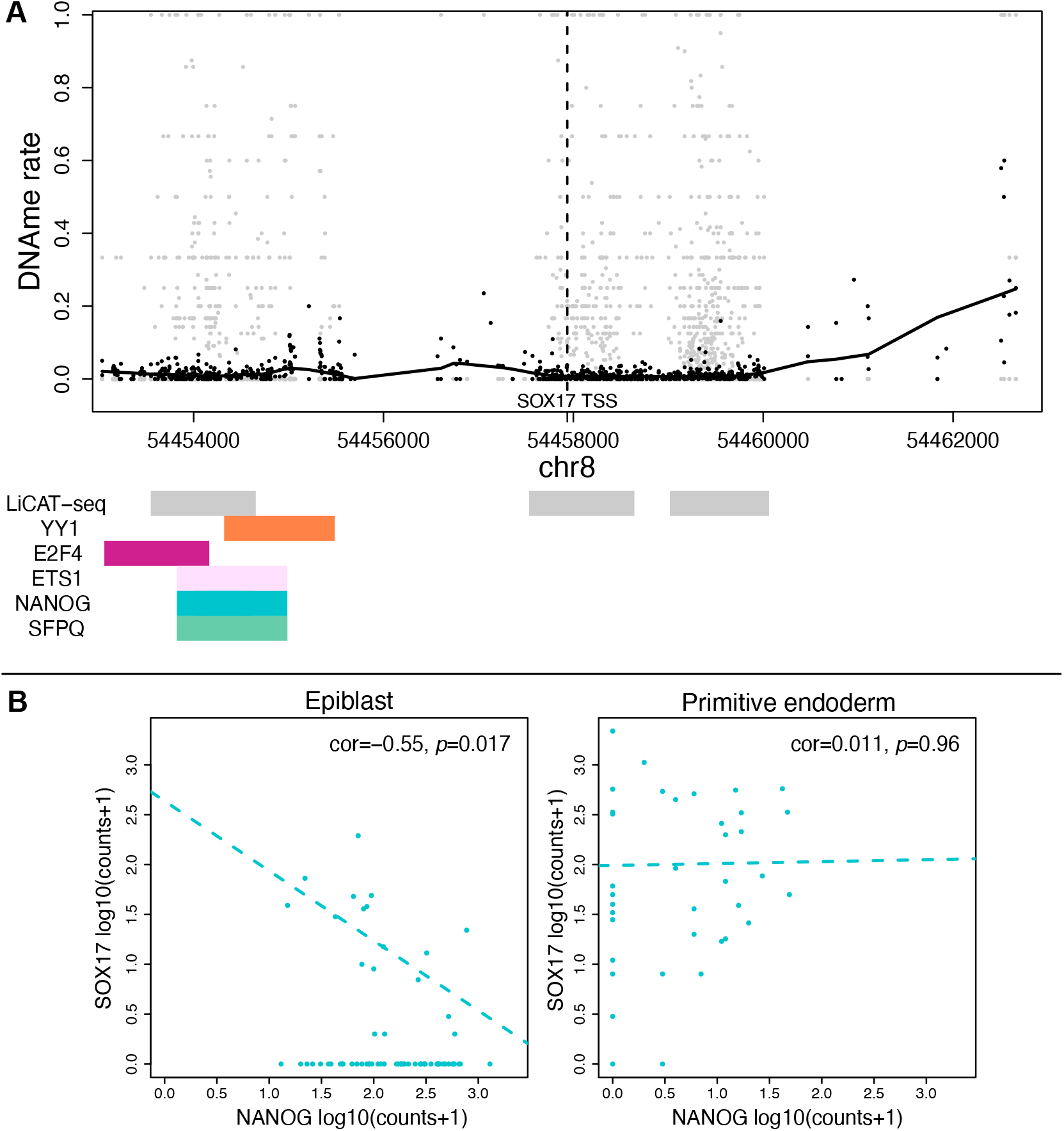
(a) The DNAme profile of the cis-regulatory region (+/-5kb) around the TSS (transcriptional start site) of the SOX17 gene (used as a marker for primitive endoderm cells in Figure S1, Figure S2, and Figure S3). Also shown are regions corresponding to previously-published and validated LiCAT-seq peaks [12, 39], and regulatory regions corresponding to motifs of candidate TF regulators inferred as part of the GRN. Mean DNAme rates are shown in black, and those for individual cells are shown in grey; non-linear trend-line is calculated with LOESS regression. (b) Scatter plots of log expression levels of SOX17 and one of its regulators NANOG [43] in Epi and PrE.

### 2.4 Differential regulation in epiblast and primitive endoderm cells

Differentially regulating TFs were identified as follows. For each TF, the differences in regulatory strength (measured by random-forests importance gain) were tested in Epi compared to PrE, for all genes the TF can regulate according to the epigenomic prior network. Figure 6a-b shows that the differential regulation in Epi and PrE is characterised by consistent upregulation of genes in PrE, whereas in Epi there is significantly more downregulation of genes (*p* < 0.001, two-sample *z*-test of proportions). This follows from the differential expression analysis (comparing Epi with PrE, Section 4), where the majority of genes were found to be upregulated in PrE, relative to Epi. We note that this is a feature of differential gene regulation that we would expect to find if the epiblast can be thought of as the default cell type at the lineage bifurcation of the epiblast and hypoblast (primitive endoderm), as has been shown recently in human embyros [46]. In this case, specification of the hypoblast / primitive endoderm should be characterised by the activation in PrE of a characteristic GRN relative to Epi, such as the differential regulation network shown in Figure 6.

**Figure 6.**
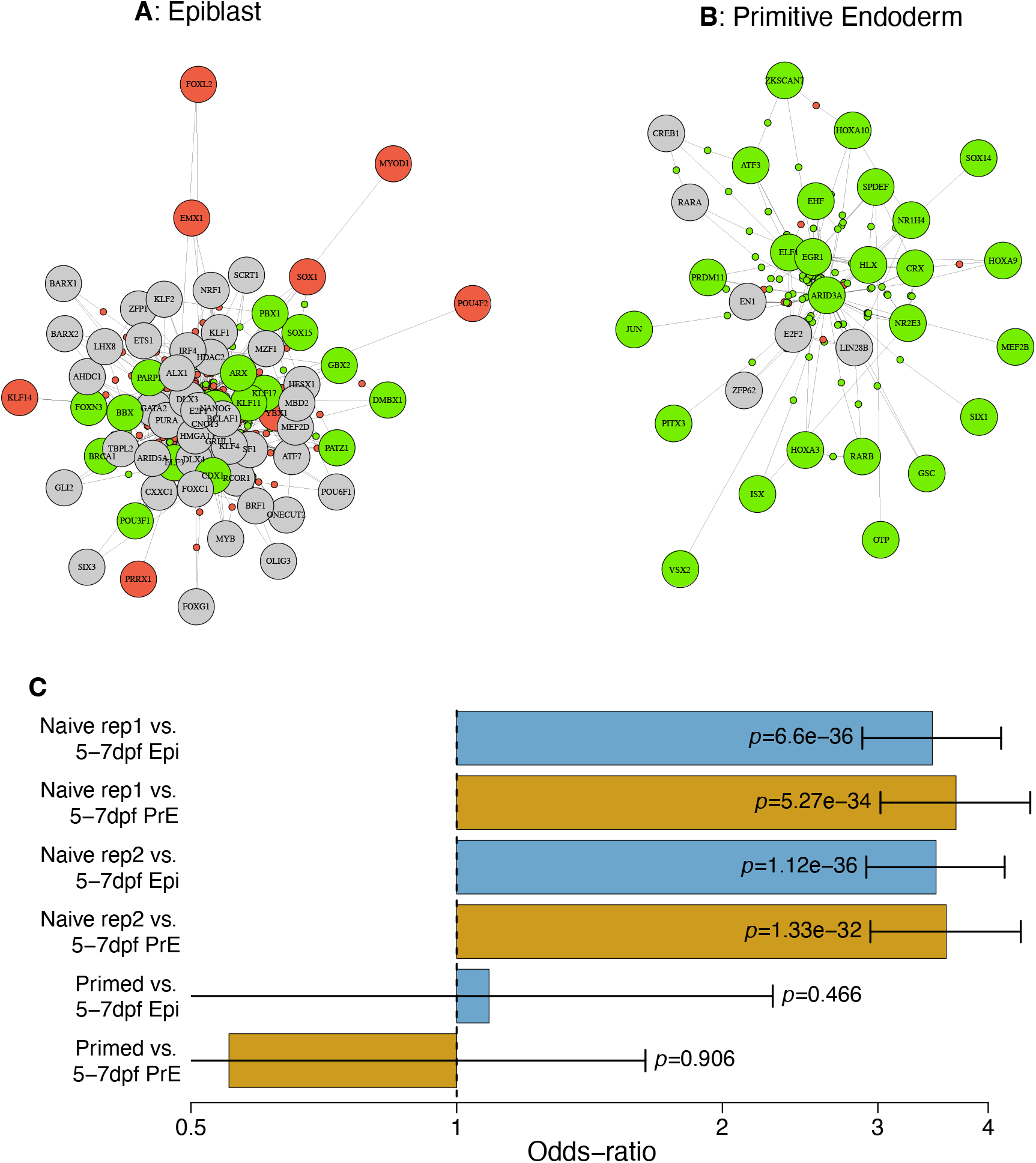
(a) Epiblast (Epi) and (b) primitive endoderm (PE) differential regulation networks. Large circles indicate significantly differentially regulating transcription factors (Table S2); green and red indicate (respectively) significantly upregulated and downregulated TFs and genes; significance level is FDR p-val < 0.05, (t-test, Benjamini-Hochberg adjustment). (c) Overlap of KLF4 targets in our GRNs inferred for 5-7dpf Epi and PrE cells, with those inferred by Cut&Tag in naive (i.e., 5-7dpf) and primed hESCs.

**Figure 7.**
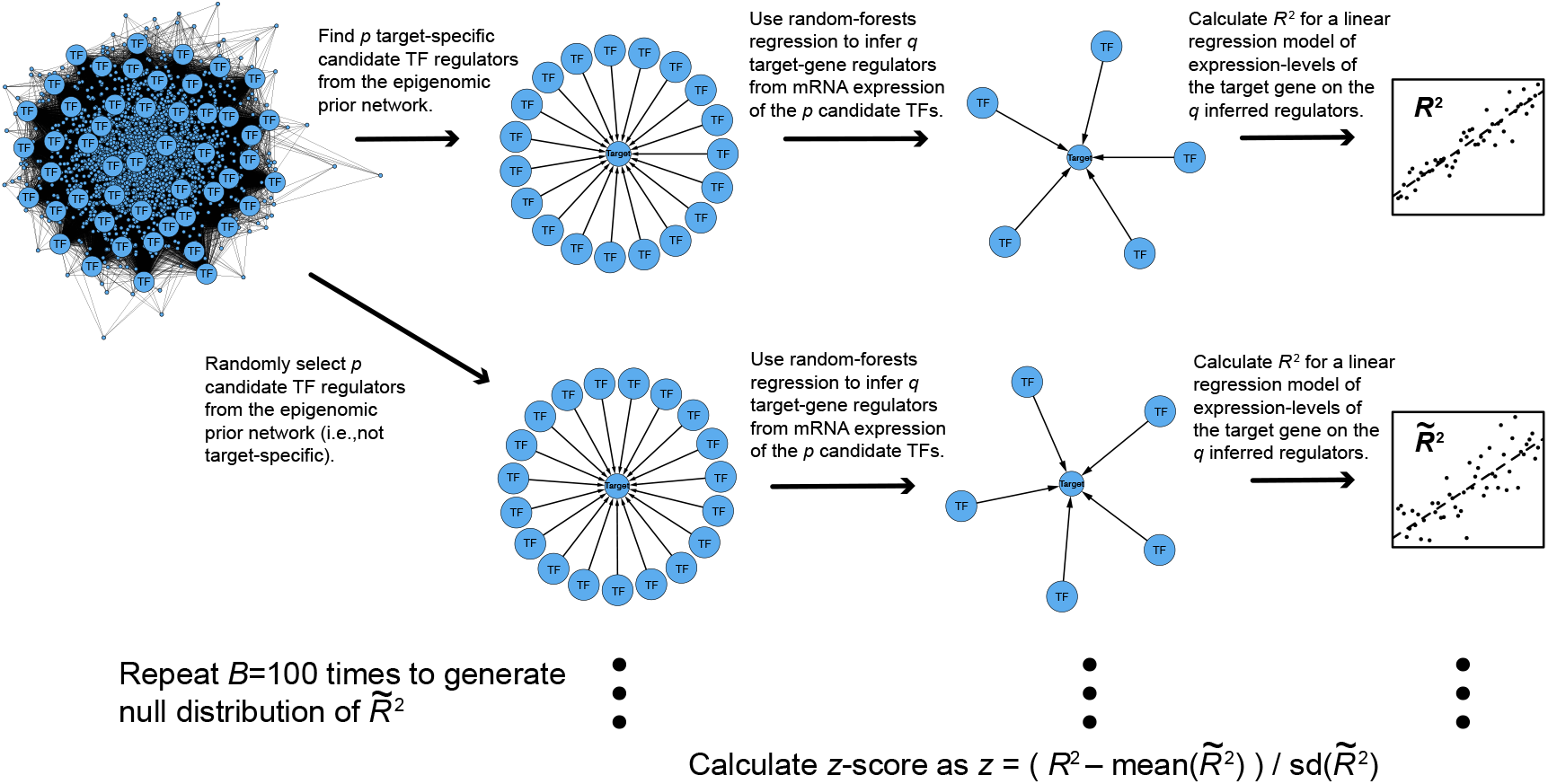
Overview of the ablation study. As in our proposed methodology (Figure 1), separately for each target gene, the epigenomic prior network defines p candidate regulators. Random-forests regression is then used to choose q < p of these p candidates, with the expression of these p TFs as predictors and the target-gene expression as the response. The R^2^ statistic quantifies the proportion of variance in the target-gene expression, that is explained by the expression of these q inferred regulators, in a linear model. A null-distribution is then generated for comparison, by randomly selecting p predictors from all those found in the epigenomic prior network (i.e. not target-specific), then applying the rest of the procedure described above to calculate a null-distribution R^2^ statistic in the same way (which we call 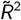). This randomisation is repeated 100 times to obtain a null-distribution of 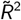. A z-score for the R^2^ statistic for this target-gene is then calculated using the mean and standard-deviation of these 100 null-distribution samples of 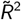.

In order to validate the targets of a specific differentially-regulating TF inferred by our proposed methodology, we re-analysed a Cut&Tag data-set of KLF4 binding to DNA, in naive and primed human embryonic stem cells (hESCs) [47]. We found that the KLF4 targets we inferred overlapped highly significantly (Fisher’s exact test) with those inferred with Cut&Tag in naive (i.e., 5-7dpf) hESCs, but not in primed hESCs (Figure 6c). We also tested enrichment of the TF targets of the differentially regulating transcription factors shown in Figure 6, by comparison with 713 epiblast gene-sets defined and validated previously by domain-experts [12]. We confirmed that there was significantly more enrichment of these epiblast gene-sets amongst TF targets differentially regulated in epiblast compared to primitive endoderm (*p* = 0.05, *t*-test), i.e., when comparing the enrichment of TF targets in Figure 6a with the enrichment of those in Figure 6b.

### 2.5 Breast cancer at-risk prior epigenomic network

Luminal progenitor, mature luminal, and basal cells are epithelial cell subtypes that are thought to have important roles in breast cancer [48], and we have previously developed and validated DNAme cancer biomarkers based on definitions of these cell types [27, 34]. We inferred epigenomic prior networks for each of the cell types luminal progenitor, mature luminal, basal, and fibroblasts, using MethCorTarget with the breast cancer at-risk DNAme dataset [49], and using MLC with the breast cancer at-risk H3K27ac ChIP-seq dataset [49]. We chose to validate the DNAme epigenomic prior network against an epigenomic prior network inferred from H3K27ac ChIP-seq data, because H3K27ac is a histone modification that is associated with activated gene-regulatory regions of the DNA [50, 51]. We found a highly significant overlap between the TF-gene regulations inferred by MLC applied to the H3K27ac data and the regulations inferred by MethCorTarget applied to the DNAme data in all cell types, hence validating the inferred DNAme epigenomic prior networks. The odds-ratios for the overlap between the gene regulations inferred from H3K27ac data and DNAme data according to these methods are OR = 45.5 (44.8 − 46.2), OR = 45.7 (45.0 − 46.5), and OR = 43.2 (42.4 − 44.0), in luminal progenitor, mature luminal, and basal cells (respectively); statistical significance is *p* < 0.001 (Fisher’s exact test) for all cell types. We also note that, interestingly, the epigenomic prior networks for luminal progenitor, mature luminal, and basal cells inferred from DNAme data again show candidate TF cis-regulations for at least eight time as many genes (respectively, 34792, 34680, and 34021 genes) than for chromatin accessibility (H3K27ac) data (respectively 4222, 4194, and 3597 different genes). Histograms showing the numbers of target-gene regulations for each TF, and the number of TF regulators for each target-gene, in the epigenomic prior networks for the breast cancer at-risk dataset, are shown in Figure S5.

### 2.6 Breast cancer at-risk GRN inference and ablation study

Women who are carriers of a mutation in the *BRCA1* (*FANCS*) or *BRCA2* (*FANCD1* genes have a greatly increased chance of suffering from breast cancer during their lifetime [52]. Hence, apparently healthy cells from *BRCA1/2* mutation carriers are likely to harbour pre-cancerous genomic changes, and insights can be gained into how breast cancer develops by comparing healthy cells from *BRCA1/2* mutation carriers (BRCAmut) with cells from *BRCA1/2* wild-type (WT). Separately for BRCAmut and WT, using the cell type specific DNAme epigenomic prior networks obtained from the DNAme breast cancer at-risk dataset [49] for luminal progenitor, mature luminal, and basal epithelial cell subtypes, as well as fibroblast cells, we inferred GRNs as described (Section 4) from the combined breast cancer at-risk RNA-seq datasets [53–59], based on the definitions of these cell types shown in Figure S4. We note that the fibroblast cells that we have labelled ‘*type 1*’ and ‘*type 2*’ are likely to correspond to (respectively) interlobular and lobular fibroblasts, based on CD26 and CD105 expression (assessed here via respectively DPP4 and ENG mRNA expression in scRNA-seq data) [60]. Lobular fibroblasts share similarities with mesenchymal stem cells; they support epithelial growth and morphogenesis, and provide a specialised microenvironment for luminal epithelial progenitors [60]. Separately for each of these epithelial and fibroblast cell-subtypes, we inferred GRNs from scRNAseq data separately for *BRCA1/2* wild-type (WT) and *BRCA1/2* mutation carriers (BRCAmut).

We carried out an ablation study (Figure 7) using the combined breast cancer at-risk dataset, to test the claim that the prior epigenomic network can significantly inform gene-regulatory network inference. We calculated *R*^2^ statistics to quantify how much of the variation in target-gene expression is explained by the TF regulators inferred for this gene target in our GRN. We then compared these observed *R*^2^ against a null distribution of *R*^2^ statistics which we call 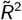, which are generated by applying our proposed GRN inference procedure to randomisations of the epigenomic prior network. We hence use the observed *R*^2^ with the mean and standard-deviation of the null-distribution of 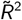 statistics to calculate a *z*-score for each target-gene. *T*-tests show that genome-wide, these *z*-scores are significantly greater than 0 (Table 1), consistently across cell-types and conditions, demonstrating that the prior network significantly informs gene-regulatory network inference.

### 2.7 Differential regulation in cells-of-origin of breast cancer

Significantly differentially regulating (diff-reg) TFs (Table S3, Table S4, Table S5, Table S6, Table S7) were identified by testing the differences in regulatory strength (measured by random-forests importance gain) in BRCAmut compared to WT, for all genes with physical evidence that cis-regulation is possible by that TF according to the epigenomic prior network. Luminal progenitor cells are hormone-receptor negative cells that are known to be the cell-of-origin of aggressive triple-negative breast cancer (TNBC) [48], and *BRCA1/2* mutation carriers are known to harbour pre-cancerous changes to in DNAme profiles [24, 26]. Hence, cis-regulatory regions of target-genes of significantly diff-reg TFs in *BRCA1/2* mutation carriers (BRCAmut) compared to wild-type (WT) in luminal progenitor cells (Figure 8a and b and Table S3) may harbour pre-cancerous alterations to DNAme profiles (methylomes). Therefore, we verified the significance of the *inferred diff-reg TFs* with respect to established knowledge, as follows. As an example, we considered the most strongly hyper-regulating TF in BRCAmut (compared to WT) in luminal progenitor cells (Table S3), the TF regulator *BACH1*; other examples can be found similarly. *BACH1* has recently been shown to be expressed more highly in TNBC [61]. Given the elevated risk of breast cancer in *BRCA1/2* mutation carriers, together with the knowledge that luminal progenitors are the cell-of-origin of TNBC, we would expect to see signs of increased regulation (hyper-regulation) by *BACH1* in luminal progenitors in *BRCA1/2* mutation carriers, as observed here. Corruption of the DNAme architecture [28] often precedes malignant transformation in cells by providing an ‘epigenetically permissible environment’, also known as an ‘epigenetic field defect’ [24, 26], and hence pre-cancerous alterations to the methylome can can serve as prognostic biomarkers [27]. Therefore, the GRN inference methodology proposed here could be used as a basis on which to develop DNAme biomarkers of cancer early-warning. Hence, we verified the significance of gene-targets of the inferred diff-reg TFs with respect to established knowledge, as follows. As an example, we looked in more detail at the gene found to be co-regulated by the largest number of diff-reg TFs in BRCAmut (compared to WT) in luminal progenitor cells, the gene *PRR7* (Figure 8); other examples can be found similarly. The *PRR7* gene is hypo-regulated in BRCAmut compared to WT, with inferred regulation by 5 and 10 diff-reg TFs, respectively (Figure 8). The *PRR7* promoter has three CpGs annotated on the Illumina 450K DNAme microarray, and there are publicly-available TCGA reference data of this type for breast cancer (BRCA) [62]. Therefore, we tested the prognostic significance of these three CpGs in the TCGA BRCA data in a survival analysis (Cox regression adjusted for confounding due to sample cell-type heterogeneity and clinical covariates), finding that the methylation level of two out of the three CpGs is significantly associated with patient survival outcome (Figure 8 and Table S8). As this gene is hypo-regulated in the at-risk population (*BRCA1/2* mutation carriers), we would expect that promoter hypermethylation at this gene would indicate poor prognosis, as observed here.

**Figure 8.**
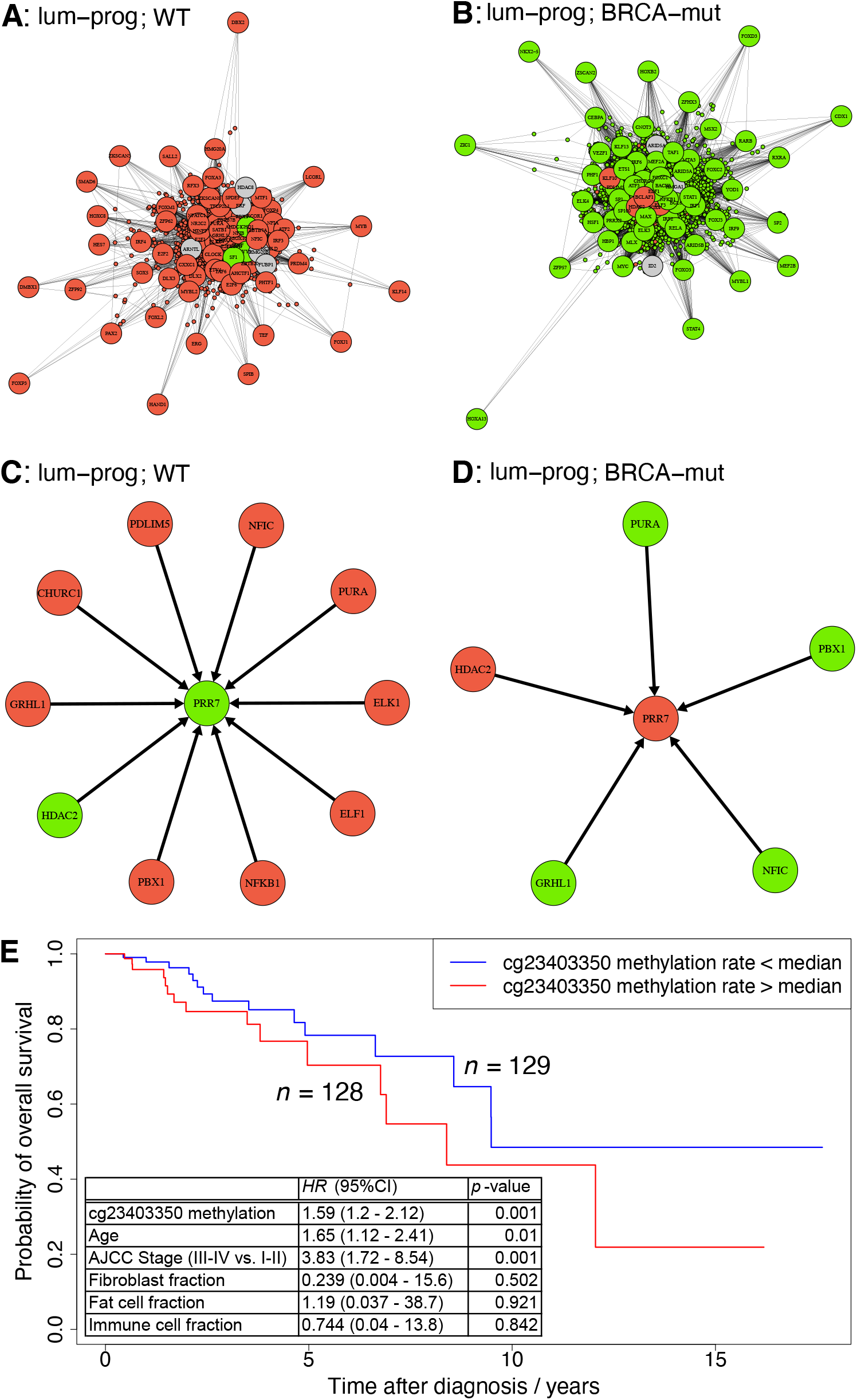
The gene-target PRR7 was found to have the largest number of regulations by differentially regulating (diff-reg) TFs of all genes, genome-wide. It is hence used as a case-study building on known, published results. (a) and (b) show the differential-regulation networks for luminal-progenitor cells for WT and BRCAmut (respectively). I.e., the part of the inferred GRNs containing differentially regulating (diff-reg) TFs (t-test, adj-p < 0.05) and differentially expressed target-genes (LIMMA, adj-p < 0.05). TFs and target-genes are shown with large and small circles respectively. (c) and (d) show the part of the GRN containing the target-gene PRR7 and its inferred diff-reg TF regulators, in WT and BRCAmut respectively. (e) Survival analysis (Cox regression adjusted for confounding by cell-type heterogeneity and clinical covariates) based on the TCGA BRCA dataset [62], for one of three CpGs (cg23403350) that are annotated to the PRR7 promoter on the Illumina 450K DNAme microarray (two of these three CpGs, cg23403350 and cg04110559, are significantly associated with survival outcome, Table S8).

In previous work [12], we tested the robustness of the inferred GRN by calculating a reproducibility or robustness score, defined as an estimate of Pr(*E*| *D*) for network edge *E* and data-set *D* [63]. In that work [12], we used the bootstrap to estimate Pr(*E*| *D*). However, we sometimes encountered computational problems using this approach with random-forests regression, due to the repeated values present in a bootstrap sample. As an alternative approach which is better suited to random-forests regression, we estimated the reproducibility or robustness score as Pr(*E* | *D*) via the Bayesian bootstrap [64, 65]. Figure 9 shows histograms of 10^5^ bootstrap samples of target-gene regression random-forests importance-gains for the regulators of *PRR*7 shown in Figure 8. The robustness or reproducibility score Pr(*E* | *D*) can then be calculated as the fraction of the 10^5^ bootstrap samples of importance-gain that are greater than the optimal threshold (Figure S6), for a given TF regulation *E* of target *PRR*7. However, not all the inferences of regulators of *PRR*7 shown in Figure 8 are found to be robust according to the analysis shown in Figure 9. This is firstly due to the high false-positive rate in network inference based on observational data [66–68], which applies equally to other state-of-the-art methods [1, 3–6, 12]. It is also likely that in some cases TF regulators of a target-gene have highly co-linear expression levels. This co-linearity would often lead to only one of these TF regulators being ‘selected’ as a regulator by advanced regression methods (such as random-forests and penalised regression) that have been found to be most effective in target-gene GRN inference [1, 3–6, 12]. As a result, true TF regulators with co-linear expression values may at random not be selected as a regulator, decreasing the corresponding estimate of Pr(*E*| *D*). Calculating bootstrap estimates of Pr(*E*| *D*) is also computationally intensive (the 10^5^ samples in Figure took 21.2 hours to generate on one core; compare with Figure S7), so it would become too computationally costly to calculate Pr(*E* | *D*) for all candidate gene-regulations, genome-wide. However, this approach can still be used for a finer-grained assessment of candidate gene-regulations, for possible further experimental investigation. Hence, for a small subset of target-genes of interest, we recommend calculating a robustness or reproducibility score according to a bootstrap estimate of Pr(*E* | *D*) as proposed by Pe’er [63].

**Figure 9.**
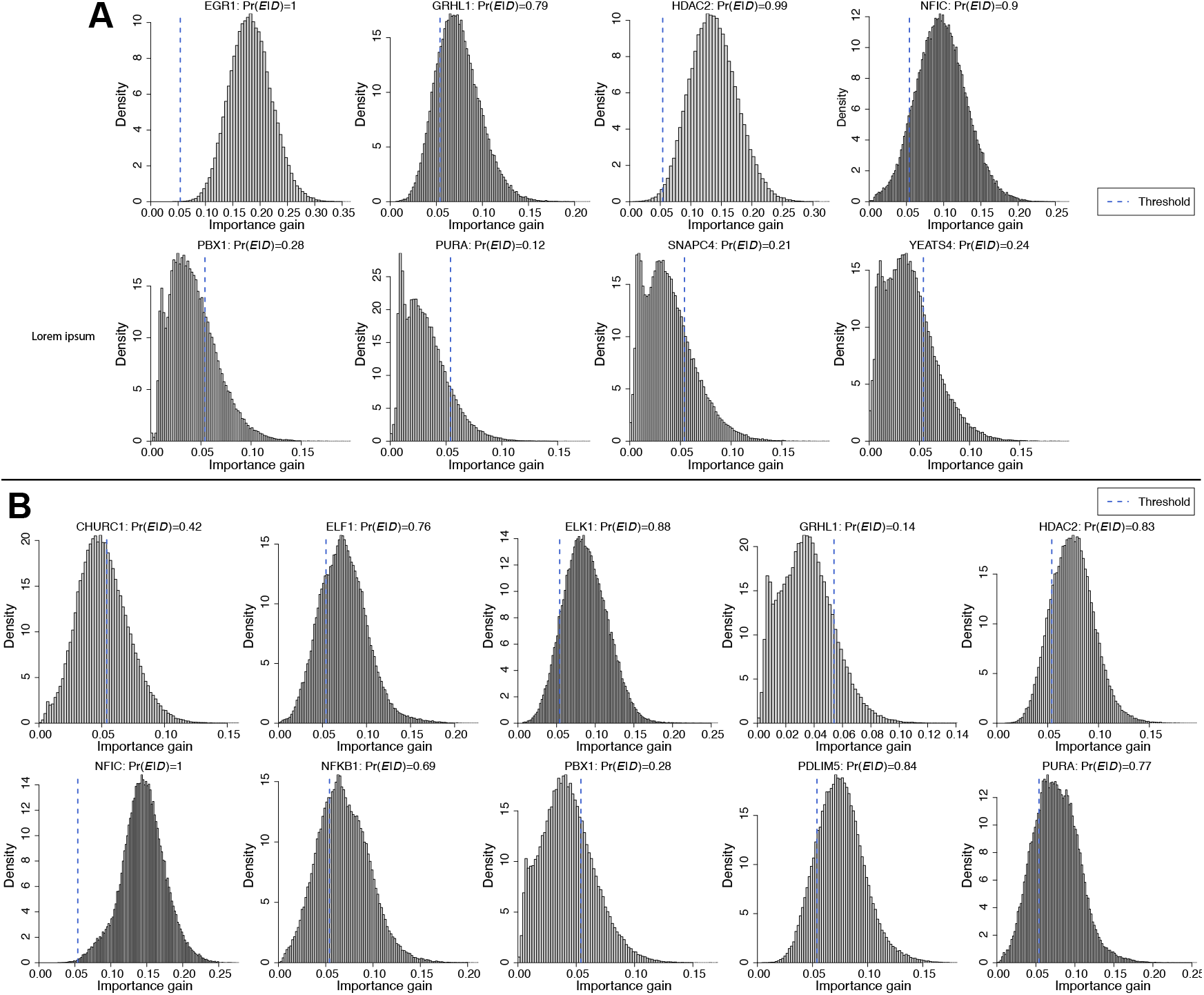
Distributions of bootstrap samples of random-forests importance-gains for the inferred regulators of PPR7 shown in Figure 8, for (a) BRCAmut and (b) WT. A GRN edge E would be inferred from data D if the importance-gain is greater than the optimal thresholds shown.

## 3 Discussion

In this work, we have shown *in-silico* that the accuracy of inference of GRN TF-gene cis-regulations is improved when epigenomic prior network structure is used in combination with advanced regression methods and gene-expression data. Furthermore, we have shown that DNAme data can be used successfully and effectively to estimate epigenomic prior network structure. There is a wealth of existing and publicly-available DNAme datasets for a wide range of tissue types that can now be leveraged using our proposed methodology to improve inference of gene-regulatory processes [13]. This contrasts with novel ATAC-seq datasets that are expensive to generate, and are currently less numerous in the public domain. However, as the availability increases of single-cell ATAC-seq data and of matched single-cell ATAC-seq and RNA-seq data, it will be interesting to see how our proposed methodology may be complemented and enhanced by the addition of single-cell chromatin accessibility data. For example, we note that single-cell ATAC-seq data are typically very sparse. Therefore, a large number of cis-regulatory regions would be expected to have low coverage in such data, and particularly in these regions the DNAme epigenomic prior network may provide useful complementary information.

We have also shown that with real data from well-studied contexts, the most significant findings from our methodology are well supported by existing literature. This gives confidence that our methodology will provide verifiable inferences in new contexts. However, we also emphasise that novel findings from computational GRN inferences remain hypotheses until they are experimentally validated. Further investigation of the differentially regulating transcription factors and their targets shown in Figure S8 and Table S3, Table S4, Table S5, Table S6, and Table S7, may allow gene-regulatory processes to be identified that are important to the oncogenic potential of these cell-types in *BRCA1* (*FANCS*) and *BRCA2* (*FANCD1*) mutation carriers, as well giving clues about possible pharmacological interventions. For example, Figure 8 shows how the methodology proposed here can be used to identify CpG loci that show altered methylation levels prior to cancer formation in at-risk tissues and populations. Hence, testing the DNAme rate of CpGs in at-risk individuals in certain cis-regulatory regions of tumour suppressor genes or oncogenes, identified by the methodology proposed here, could provide important early-warning biomarkers of cancer or of cancer predisposition. However, further work will be required to systematically find and then validate all the examples of changes in DNAme patterns that may be discovered with our proposed methodology, and that are associated with changes in TF cis-regulation of specific genes that precede the onset of disease.

Previous advanced methods for GRN inference, such as GENIE3 [5], inferred GRNs based only on gene expression data. By including TF motif-DNA matches along with epigenomic data such as DNAme data, we make our GRN inferences more physically realistic. However, chromatin state (such as histone modifications), as well as the interaction of co-factors, and sequence context, all play an important part in determining gene regulation. Therefore, future development of our methodology will incorporate a wider range of ‘omics and sequence data to further refine the GRN inferences. While we have shown that our proposed methodology is effective in the context of datasets of a wide range of sizes from the very different contexts of human embryonic development, healthy individuals at-risk of breast cancer, and individuals with advanced breast cancer, we acknowledge that this methodology remains untested outside of these contexts. Future work planned, includes expanding our proposed methodology in a tailored way to a wide range of other applications. This will include making a wide range of epigenomic prior networks available for different cell-types.

## 4 Methods

### 4.1 Quantification and statistical analysis

#### 4.1.1 Epigenomic prior network construction

To obtain the candidate TF cis-regulators of a particular gene, we build on established methods called lever [14] and cistarget [15]. Those methods examine whether the association between TF binding motifs and cis-regulatory DNA sequences corresponds to the association between sample-specific observations and the same cis-regulatory DNA regions, such as open chromatin (as inferred from ATAC-seq or LiCAT-seq data). Lever [14] and cistarget [15] approach this as a prediction problem: i.e., does the matching of TF motif and DNA sequence at a particular location predict epigenomic patterns (such as open chromatin) that are observed at that location. In lever [14] and cistarget [15], the accuracy of this prediction is measured by the AUC (area under receiver-operator curve) statistic. The AUC statistic can be calculated from the area under the curve that plots the true-positive rate against the false-positive rate, as the classification threshold is decreased from infinity to zero; i.e., the receiver-operator curve. Equivalently, the AUC statistic can be calculated by a linear transformation of Somers’ *D* statistic by the transformation AUC = (*D* + 1)*/*2 [69], where *D* ∈ [−1, 1]. Somers’ *D* statistic can also be thought of as equivalent to a correlation coefficient between a continuous variable (the prediction) and a binary variable (the ground-truth). If the correlation coefficient *ρ* ∈ [−1, 1] is a measure of association between two binary variables, then *D* ∈ [−1, 1] can be thought of as an analogous measure of association between a binary variable and a continuous variable. From the perspective of the AUC statistic, the binary variable corresponds to a representation of the true class *y* ^(*r*)^ ∈ {0, 1} for rankings *r* ∈ {1, …, *R*}, and the continuous variable *x* ^(*r*)^ is used to predict that class, where a value of the AUC statistic close to 1 indicates that high rankings mostly correspond to a true class of 1.

In MethCorTarget, we use the absolute Spearman correlation coefficient to assess whether the average DNAme rate 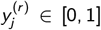 for sample *j* ∈ {1, …, *n*} at cis-regulatory region *r* ∈ {1, …, *R*} corresponds to the closeness of the match 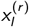 between TF motif *l* ∈ {1, …*L*} and the DNA sequence at that region, where correlation coefficient 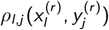 is calculated over *r* ∈ {1, …, *R*}. In modified lever-cistarget (MLC), we equivalently use Somers’ *D* to assess whether the presence of open chromatin (assessed by LiCAT-seq or H3K27ac) at a cis-regulatory region corresponds to the closeness of the match between the TF motif and the DNA sequence at that region. After calculating *ρ*_*l,j*_ (for a given DNAme sample *j*) or *D*_*l,j*_ (for other epigenomic data sample *j*) for TF motif *l*, we apply the Fisher transform to obtain approximately standard Gaussian-distributed variables *z*_*l,j*_ ~ 𝒩 (0, 1) [70], *l* ∈{1, …*L*}, *j* ∈{1, …, *n*}, for *L* TFs and *n* samples (assuming the distributional assumptions also hold sufficiently well for *D*), and then calculate corresponding *p*-values from a standard Gaussian distribution. We then retain TFs in the subsequent analysis if the corresponding *z*-test leads to *p*-value *p*_*l,j*_ < 0.1, noting that this threshold provides optimum identifiability of network structures [71]. We also note that this *p*-value threshold might result in many false-positives for a large number of TFs, but that many of these would be expected to be filtered out in the next step in the GRN inference using gene expression data with advanced regression methods. The aim of the epigenomic prior network is to identify candidate regulators of a target-gene, which are then chosen from by the advanced regression method as it is applied to gene-expression data. We refer to this method of epigenomic prior network inference as MLC (modified-lever-cistarget) when the method calculates Somers’ *D* from chromatin accessibility data (or similar), and we refer to this method as MethCorTarget when the method calculates the Spearman correlation coefficient *ρ* from DNAme data.

Histograms showing the numbers of target-gene regulations for each TF, and the number of TF regulators for each target-gene, in the epigenomic prior networks for the breast cancer at-risk dataset, and for the human embryonic development dataset, are shown in Figure S5.

#### 4.1.2 Gene-regulatory network inference

Gene-regulatory network (GRN) inference was carried out to infer TF-gene regulations by modelling the expression of each target-gene as a (possibly non-linear) combination of the expression of a set of candidate regulators (Equation 1) [4]. The target-gene approach to GRN inference [4] uses advanced regression methods such as random forests regression [72], and recently the xgboost implementation of random forests as gradient-boosted trees [38], to model the expression level of a particular ‘target’ gene (the response), conditional on the expression levels of a set of other genes (predictors, such as TFs), Equation 1. The target-gene approach to GRN inference is best known in the context of GENIE3 [5] and GRNboost2 [6]. In GENIE3 and GRNboost2, the set of candidate regulators corresponds to all the TFs in the dataset of interest, or to some other pre-specified list of candidate regulators that importantly is the same for all target-genes. In our proposed methodology, we use the epigenomic prior network to first infer a specific set of candidate TF regulators that can be different for each gene (Figure 1). This improves the accuracy of inference by the advanced regression method (Figure 3), because when using advanced regression such as random forests for variable selection, the advanced regression only needs to choose from 10s of candidate cis-regulating TFs. These 10s of candidate cis-regulators correspond to epigenomic prior information about possible interactions of the TF with the cis-regulatory DNA of the target-gene. This contrasts with the full list of 100s or 1000s of TFs that would need to be specified genome-wide when using GENIE3 [5] and GRNboost2 [6], as part of the SCENIC / SCENIC+ pipeline [1–3]. Finally, we construct the GRN by combining the local target-gene network model fits genome-wide, with each gene taking a turn as the response variable in the advanced regression, i.e., as the target-gene [4]. In the final GRN, we infer network edges using optimised thresholds on the random-forests importance-gain in each target-gene model fit. We estimated optimal thresholds based on a maximum-AUC criterion in data with known ground-truth network structure, for relevant values of sample size *n* and number of parents in the network *p* (Figure S6). The practitioner is free to choose the threshold as a user parameter, based on considerations of the downstream analyses. We choose to set the threshold as the lower-quartile of the optimal AUC value conditional on sample size *n* and number of parents in the network *p*. This lower choice of threshold ensures that false-negatives are reduced at the price of additional false-positives, which is preferred for two reasons. Firstly, further downstream validation will be used, mitigating any increase in false-positives. Secondly, downstream we analyse aggregations of gene regulations, which also makes the downstream analyses more tolerant to false-negatives, as their effect is mitigated when aggregated with true positives (Figure S6).

We found previously [12] that advanced regression methods as part of the target-gene approach to GRN inference perform best with normalisation methods such as FPKM and TPM. In this work, we chose to use CPM normalisation rather than FPKM or TPM normalisation, because inferences from advanced regression with the random forests method should not be affected by monotonic transformations of the data, such as scaling by gene length. In comparison with that previous work [12], we also chose to use random forests regression [72] (xgboost implementation [38]) rather than mutual information, for three reasons as follows: (1) The computational efficiency of xgboost [38]) makes it feasible to use xgboost with large, modern single-cell datasets such as the breast cancer at-risk dataset we analyse in this work. This contrasts with the much smaller number of cells in the human embryonic development dataset, where cell numbers are restricted due to the very precious and restricted nature of the source material; (2) In our previous work [12] we used the GENIE3 implementation [5] of target-gene GRN inference with random-forests [72], but in this work we have have shown a clear improvement over GENIE3 on synthetic data by using prior network structure (Figure 3), noting that GENIE3 does not use prior network structure that is tailored to each target-gene; (3) In this work we have based our epigenomic prior network inference methods on the lever [14] / cistarget [15] approach, which is also used together with random forests [72] as part of the well validated scenic [1, 3] pipeline. On the other hand our previous work [12] used RGT-HINT [36] to infer candidate TF-gene cis-regulations.

#### 4.1.3 LiCAT-seq data processing

For the human embryonic development LiCAT-seq dataset [39], the DNA loci of cis-regulatory TF motifs corresponding to open chromatin were inferred as part of an earlier study [12] (with alignment to the GRCh38 reference human genome), and were reused in this work. These TF motif footprints at open chromatin were originally inferred from the human embryonic development LiCAT-seq dataset available from the NCBI Sequence Read Archive (SRA) under accession number SRP163205, from the URL https://www.ncbi.nlm.nih.gov/sra/?term=SRP163205 The TF-motif footprints at open chromatin were originally obtained using rgt-hint v0.13.0 [36] with TF binding models from HOCOMOCO and JASPAR [39], finding 433270 candidate cis-regulations identified over 13950 different genes by 894 different TFs.

We applied pycistarget and modified-lever-cistarget (MLC) to the downstream peak-calls (54425 peak-calls genome wide) for the human embryonic development LiCAT-seq dataset [39] for comparison. When applied to these peak-calls, pycistarget returned 124 significant TF motifs across 39298 distinct genomic regions. The MLC method found 549 significant TF motifs, matched across 27280 distinct genomic regions (*z*-test, *p* < 0.1, we note that many false positives generated in this step are expected to be removed by the subsequent GRN inference with gene-expression data). For MLC, candidate cis-regulations at these 27280 genomic regions were inferred from the closest gene promoters (< 5kb, according to Ensemble 107 [73]). This lead to a epigenomic prior network for the human embryonic development LiCAT-seq dataset, representing 21381 candidate cis-regulations of 2650 different genes by 270 different TFs.

#### 4.1.4 H3K27ac ChIP-seq data processing

For the breast cancer at-risk H3K27ac ChIP-seq dataset [49], raw fastq files of unaligned sequencer reads for purified cell samples for the cell types luminal progenitor, mature luminal, and basal from each of 3 women (aged 22, 37 and 41), were downloaded from EGA (the European Genome-phenome Archive, https://ega-archive.org), under accession number EGAS00001000552. ChIP-seq reads were aligned to the GRCh38 (hg38) reference human genome assembly from Ensembl 109 [74] using the Burrows-Wheeler Alignment (BWA) software [75] (with default settings). Significant ChIP-seq peaks (FDR-adj *p* < 0.05) were then called using the callpeak function from the macs2 software [76, 77] (with default settings), resulting in three sets of peak calls per cell type. Applying MLC to these data for the cell types luminal progenitor, mature luminal, and basal, lead to (respectively) 583, 591, and 595 significant TF motifs matched across 25706, 27668, and 22570 genomic regions (*z*-test, *p* < 0.1, we note that many false positives generated in this step are expected to be removed by the subsequent GRN inference with gene-expression data). For each genomic region, candidate cis-regulations were inferred from the closest gene promoters (< 5kb, according to Ensemble 107 [73]), also requiring that the candidate cis-regulation should be observed at this significance level in a majority of the samples for that cell type. This lead to a epigenomic prior network for each of the cell types luminal progenitor, mature luminal, and basal, representing (respectively) 142766, 142237, and 110513 candidate cis-regulations of 4222, 4194, and 3597 different genes, by 282, 285, and 277 different TFs.

#### 4.1.5 DNAme bisulphite-seq data processing

For the *human embryonic development* DNAme dataset [40], CpG read-count data tables summarising bisulphite-seq libraries (aligned to the hg19 reference human genome) for 103 human ICM cell samples (inner cell mass, comprising epiblast and primitive endoderm cells, one library per cell sample), were downloaded from from the Gene Expression Omnibus (GEO) under accession number GSE81233, from the URL: https://www.ncbi.nlm.nih.gov/geo/query/acc.cgi?acc=GSE81233

For the *breast cancer at-risk* DNAme dataset [49], bisulphite-seq DNAme data for purified cell samples donated by 3 women (aged 22, 37 and 41, one library per cell type for each woman) for luminal progenitor, mature luminal, basal, and stromal (fibroblast) cells, were downloaded from EGA (the European Genome-phenome Archive, https://ega-archive.org), under accession number EGAS00001000552. The downloaded raw sequencer reads were aligned to the GRCh37 (hg19) reference human genome assembly from Ensembl 104 [78], and methylated and unmethylated reads were counted at CpG sites, using the Bismark software [79] (with default settings).

To generate the epigenomic prior networks from these DNAme datasets, MethCorTarget was applied to the raw methylated and unmethylated bisulphite-seq counts, as follows. First, the methylated and unmethylated counts were aggregated over all candidate TF-motif binding regions that were mapped by at least 10 reads in total, leading to an average DNAme rate for each candidate TF-motif binding region, for each sample (i.e., bisulphite-seq library). The average methylation rates over the samples were then calculated for all TF-motif binding regions, for only those regions with data for at least 3 cells in the human embryonic development dataset, and for only those regions with data for the majority of samples in the breast cancer at-risk dataset. We note that DNAme data are typically quite noisy, but by averaging the methylation rates over TF binding regions, we mitigate some of the effect of this noise, as shown previously [80].

We used MethCorTarget to test the correlation of the ranking of the TF-motif DNA sequence matches for each binding region, with the average methylation rates over the binding regions calculated as above. We applied MethCorTarget in the same way for ICM cells of the human embryonic development dataset, and for the cell types luminal progenitor, mature luminal, basal, and stromal (fibroblast) cells of the breast cancer at-risk dataset. For ICM cells, MethCorTarget found 868 significant TF motifs matched across 95122 genomic regions. For luminal progenitor, mature luminal, basal, and stromal (fibroblast) cells, MethCorTarget found (respectively), 826, 780, 747, and 771 significant TF motifs matched across 206076, 201359, 195211, and 200102 genomic regions (*z*-test, *p* < 0.1; we note that many false positives generated in this step are expected to be removed by the subsequent GRN inference with gene-expression data). For each genomic region, candidate cis-regulations were inferred from the closest gene promoters (< 5kb) to the motif-matched TFs (according to Ensemble 107 [73]). This lead to a epigenomic prior network for ICM representing 417320 candidate cis-regulations of 23575 genes by 375 TFs. It also lead to a epigenomic prior network for each of the cell types luminal progenitor, mature luminal, basal, and stromal (fibroblast) representing (respectively) 1035073, 946625, 920222, and 991998 candidate cis-regulations of 34792, 34680, 34021, and 34623 mRNA transcripts, by 413, 370, 382, and 387 TFs.

For plotting the bisulphite-seq reads aligned to hg19 on hg38 (GRCh38) co-ordinates, the liftover package in R was used to convert CpG loci from the hg19 coordinates to hg38. We note that comparisons between LiCAT-seq data aligned to hg38 co-ordinates and DNAme data aligned to hg19 co-ordinates were at the level of unique identifiers for cis-regulatory regions in the TF-motif DNA-binding database, mitigating alignment errors and loss of data that could arise during coordinate conversion.

#### 4.1.6 RNA-seq data processing

The *human embryonic development* RNA-seq dataset [42] was downloaded from ArrayExpress under accession number EMTAB3929, from URL https://www.ebi.ac.uk/biostudies/arrayexpress/studies/E-MTAB-3929

After quality control and filtering to retain cells from embryos at 5-7 days post-fertilisation, data for 20407 features (RNA transcripts) in 1258 samples (cells) were identified for use in downstream GRN inference.

The breast cancer at-risk RNA-seq datasets [53–59] have been analysed together in combination previously [58], and hence are referred to here as the *combined breast-cancer at-risk* dataset, and were downloaded from (respectively) the Gene Expression Omnibus (GEO) under accession numbers GSE161529, GSE174588, GSE195665, GSE198732, GSE180878, and ArrayExpress under accession numbers EMTAB13664, EMTAB9841, from (respectively) URLs https://www.ncbi.nlm.nih.gov/geo/query/acc.cgi?acc=GSE161529 https://www.ncbi.nlm.nih.gov/geo/query/acc.cgi?acc=GSE174588 https://www.ncbi.nlm.nih.gov/geo/query/acc.cgi?acc=GSE195665 https://www.ncbi.nlm.nih.gov/geo/query/acc.cgi?acc=GSE198732 https://www.ncbi.nlm.nih.gov/geo/query/acc.cgi?acc=GSE180878 https://www.ebi.ac.uk/biostudies/arrayexpress/studies/E-MTAB-13664 https://www.ebi.ac.uk/biostudies/arrayexpress/studies/E-MTAB-9841

After quality control and filtering to retain data from healthy pre-menopausal nulliparous women, data for 19995 features (RNA transcripts) in 70725, 65786, and 57906 samples (cells) from (respectively) 11 *BRCA1* (*FANCS*) mutation carriers, from 5 *BRCA2* (*FANCD1*) mutation carriers, and from 18 *BRCA1* (*FANCS*) and *BRCA2* (*FANCD1*) wild-type volunteers, were identified for use in downstream GRN inference.

Quality control was carried out as follows, to retain only high-quality samples (cells) and features (RNA transcripts). For both the human embryonic development dataset and the combined breast cancer at-risk dataset, samples (cells) were retained by filtering out any cells with non-zero counts for fewer than 750 RNA transcripts, filtering out cells for which the most highly expressed transcript excluding MALAT1 account for at least 10% of the total counts, filtering out cells for which at least 10% of the total counts correspond to transcripts encoding mitochondrial proteins, filtering out cells for which at least 50% of the total counts correspond to transcripts encoding ribosomal proteins, and filtering out doublets [81]. For the human embryonic development dataset, features were filtered out if they have non-zero counts in fewer than 10 samples (cells), noting the smaller sample size and greater read-depth for this dataset. For the combined breast-cancer at-risk dataset, RNA transcripts were filtered out if they have non-zero counts in fewer than 100 cells. For the combined breast cancer at-risk dataset, the COMBAT software [82] was used to control batch effects while combining datasets.

After quality control, differential expression analysis was carried out using the limma software [83] from the edgeR package [84] in R, with significance level set according to false discovery rate-adjusted *p* < 0.05. Separately after quality control, hypervariable mRNA transcripts were identified for dimensional reduction, visualisation, and clustering, following the principle of selecting genes with the greatest biological variability [85]. We use an estimator of the amount of biological variability compared to technical variability (i.e., overdispersion [85]) in each mRNA feature *i* ∈{1, …, *p*}, defined here as log*V*_*i*_ */*log*m*_*i*_ where *V*_*i*_ and *m*_*i*_ are (respectively) the empirical variance and mean for feature *i*. We note that there is currently no consensus on the best-practice for selecting the most informative features using models of overdispersion, and that the best solution is likely to depend both on the down-stream processing and on as the datasets that are analysed [86–89]. The top 2000 mRNA features were retained in this way for the breast cancer at-risk dataset, and the top 5000 mRNA features were retained for the embryonic development dataset (recognising that more features are needed for estimating the Laplacian eigenspace as this dataset contains many fewer cell samples).

The Laplacian eigenspace (LE) is the basis for the visualisations and clustering shown in Figure S1, Figure S2, and Figure S4, and is estimated as the subspace of the data defined by the eigendecomposition of the graph-Laplacian representation of the datapoints [90], according to the eigenvectors that correspond to the eigenvalues that explain at least 1% of the total variance of the data, after removing the mean component. The single-cell data for the specific cell types analysed in the GRN inference are identified from the LE with unsupervised learning by fitting a Gaussian mixture model in the Laplacian eigenspace (GMM-LE clustering) [90, 91], as shown in UMAP projections [92] from the LE [90] that appear in Figure S1, Figure S2, and Figure S4.

For the human embryonic development dataset, the epiblast cells used in the GRN inference are the same ones that we used in previous work [90], identified as cluster 5 in Figure S1; the identifiers for these Epi cells from the original study that generated those data [42] are listed in Table S9. GMM-LE clustering was then applied hierarchically to cluster 2 from Figure S1, resulting in 37 primitive endoderm (PE) cells identified as cluster 4 in Figure S2; the identifiers for these PrE cells from the original study that generated these data [42] are listed in Table S10. The 68 and 37 Epi and PrE cells (respectively) are validated in Figure S3 by the expression levels of known Epi marker genes *NANOG* and *KLF17* and PrE marker genes *GATA4* and *SOX17* [12], and are used as representative samples of gene-expression profiles of those cell types in the GRN inference. For the breast cancer at-risk dataset, we previously identified 5531 luminal progenitor, 2535 mature luminal, and 3913 basal cells [90], which are identified as clusters 1, 2, and 8, respectively in Figure S4. These cell samples are then used as representative examples of gene-expression profiles of those cell types in the GRN inference.

#### 4.1.7 DNAme bulk microarray data processing

Bulk-tissue Illumina 450K DNAme microarray data for 257 breast cancer invasive carcinoma (BRCA) samples with matched clinical data were downloaded from the TCGA repository (The Cancer Genome Atlas) [62], and were preprocessed as follows. Raw DNAme idat data files were loaded in R and background-corrected and normalised with BMIQ [93]. Probes were removed if they had < 95% coverage across samples, and any remaining probes with detection *p*-value *>* 0.05 were replaced by *k*-NN imputation, with *k* = 5.

#### 4.1.8 Cut&Tag dataset

Previously-published [47] Cut&Tag KLF4-target peak-calls (*p* < 0.01) were downloaded for naive (two replicates) and primed (one replicate) human embryonic stem cells (hESCs), from the Gene Expression Omnibus (GEO) under accession number GSE167988.

#### 4.1.9 TF-motif databases

The rankings for the matches of TF-motif to DNA sequence location for human, that we used with pycistarget, MLC (modified-lever-cistarget) and MethCorTarget, are based on 32765 unique TF motifs that were originally collated from 29 different motif databases as part of the SCENIC+ package [3], and were downloaded under the filenames `hg38 screen v10 clust.regions vs motifs.rankings.feather` (for the motif rankings) and `motifs-v10nr clust-nr.mgi-m0.001-o0.0.tbl` (for the TF-motif matches), from the URL: https://resources.aertslab.org/cistarget/

#### 4.1.10 Synthetic data generation

Synthetic gene expression data was generated, corresponding to known ground-truth GRN structure. These data were generated using the ‘gene netweaver’ software [94], which uses realistic differential-equations models to simulate the data according to models of chemical reaction kinetics; this software was used previously to generate data for the DREAM4 and DREAM5 network inference challenges [37] that were originally won by GENIE3 [5]. Using gene-net weaver, we generated 300 synthetic gene-expression profiles based on known GRN structure. These 300 synthetic gene-expression profiles were then analysed equivalently to 300 normalised single-cell libraries sequenced at high depth. The number of synthetic data samples, the maximum synthetic network size, and the number of candidate regulators of each target gene, were determined by the constraints of the data generating process

#### 4.1.11 Deconvolution of bulk DNA methylation data

Sample fractions of epithelial cells, fibroblasts, immune cells and fat cells were estimated for the TCGA BRCA bulk DNAme microarray data using the ‘Epidish’ package in R [33]. We note that only the fractions of fibroblasts, immune cells and fat cells were used to adjust for sample cell-type fraction due to built-in redundancy in these estimates (i.e., the epithelial, fibroblast, immune, and fat fractions sum to 1 by definition).

#### 4.1.12 Software

Downstream analyses were carried out in R (versions 4.1-4.3), except for the application of pycistarget to the LiCAT-seq data, which used Python version 3.10 and pycistarget version 1.0.

### 4.2 Key resources

The human embryonic development DNAme dataset is available from the Gene Expression Omnibus (GEO) under accession number GSE81233, from the URL https://www.ncbi.nlm.nih.gov/geo/query/acc.cgi?acc=GSE81233

The human embryonic development LiCAT-seq dataset is available from the NCBI Sequence Read Archive (SRA) under accession number SRP163205, from the URL https://www.ncbi.nlm.nih.gov/sra/?term=SRP163205

The breast cancer at-risk DNAme (bisulphite-seq), and H3K27ac (ChIP-seq) datasets, are available from EGA under accession number EGAS00001000552, from the URL https://ega-archive.org

The human embryonic development single-cell RNA-seq dataset is available from ArrayExpress under accession number E-MTAB-3929, from the URL https://www.ebi.ac.uk/biostudies/arrayexpress/studies/E-MTAB-3929

The breast cancer at-risk single-cell RNA-seq datasets are available from the Gene Expression Omnibus (GEO) under accession numbers GSE161529, GSE174588, GSE195665, GSE198732, GSE180878, and ArrayExpress under accession numbers EMTAB13664, EMTAB9841, from URLs https://www.ncbi.nlm.nih.gov/geo/query/acc.cgi?acc=GSE161529 https://www.ncbi.nlm.nih.gov/geo/query/acc.cgi?acc=GSE174588 https://www.ncbi.nlm.nih.gov/geo/query/acc.cgi?acc=GSE195665 https://www.ncbi.nlm.nih.gov/geo/query/acc.cgi?acc=GSE198732 https://www.ncbi.nlm.nih.gov/geo/query/acc.cgi?acc=GSE180878 https://www.ebi.ac.uk/biostudies/arrayexpress/studies/E-MTAB-13664 https://www.ebi.ac.uk/biostudies/arrayexpress/studies/E-MTAB-9841

The breast cancer invasive carcinoma (BRCA) DNAme dataset and associated demographic information is available from TCGA [62] from the URL http://www.cbioportal.org/study/summary?id=brca_tcga

The Cut&Tag KLF4 binding in hESCs dataset is available from the URL https://www.ncbi.nlm.nih.gov/geo/query/acc.cgi?acc=GSE167988

The TF-motif DNA-sequence match rankings are available from the URL: https://resources.aertslab.org/cistarget/

R scripts implementing the epigenomic prior network construction, GRN inference, and differential cis-regulatory analysis, with required data-files for the human embryonic development datasets are available from https://zenodo.org/records/17253602, and with required data-files for the breast cancer at-risk datasets from https://zenodo.org/records/17255982

## 5 Additional information

### Author contributions

TB conceived of and designed the study. TB wrote the manuscript, with advice from QH. Statistical analyses and model development were carried out by TB, with contributions from ML, CS, YG, and QH. All co-authors approved this version of this manuscript.

## Acknowledgements

The results published here are in whole or part based upon data generated by The Canadian Epigenetics, Epigenomics, Environment and Health Research Consortium (CEEHRC) initiative funded by the Canadian Institutes of Health Research (CIHR), Genome BC, and Genome Quebec. Information about CEEHRC and the participating investigators and institutions can be found at http://www.cihr-irsc.gc.ca/e/43734.html

The authors acknowledge the use of the UCL High Performance Computing Facilities, and associated support services, in the completion of this work.

## License

This manuscript is available under a Creative Commons CC BY-NC License, as described at https://creativecommons.org/licenses/by-nc/4.0/

