## Supplementary figures and tables for "Inferring gene-regulatory networks using epigenomic priors"

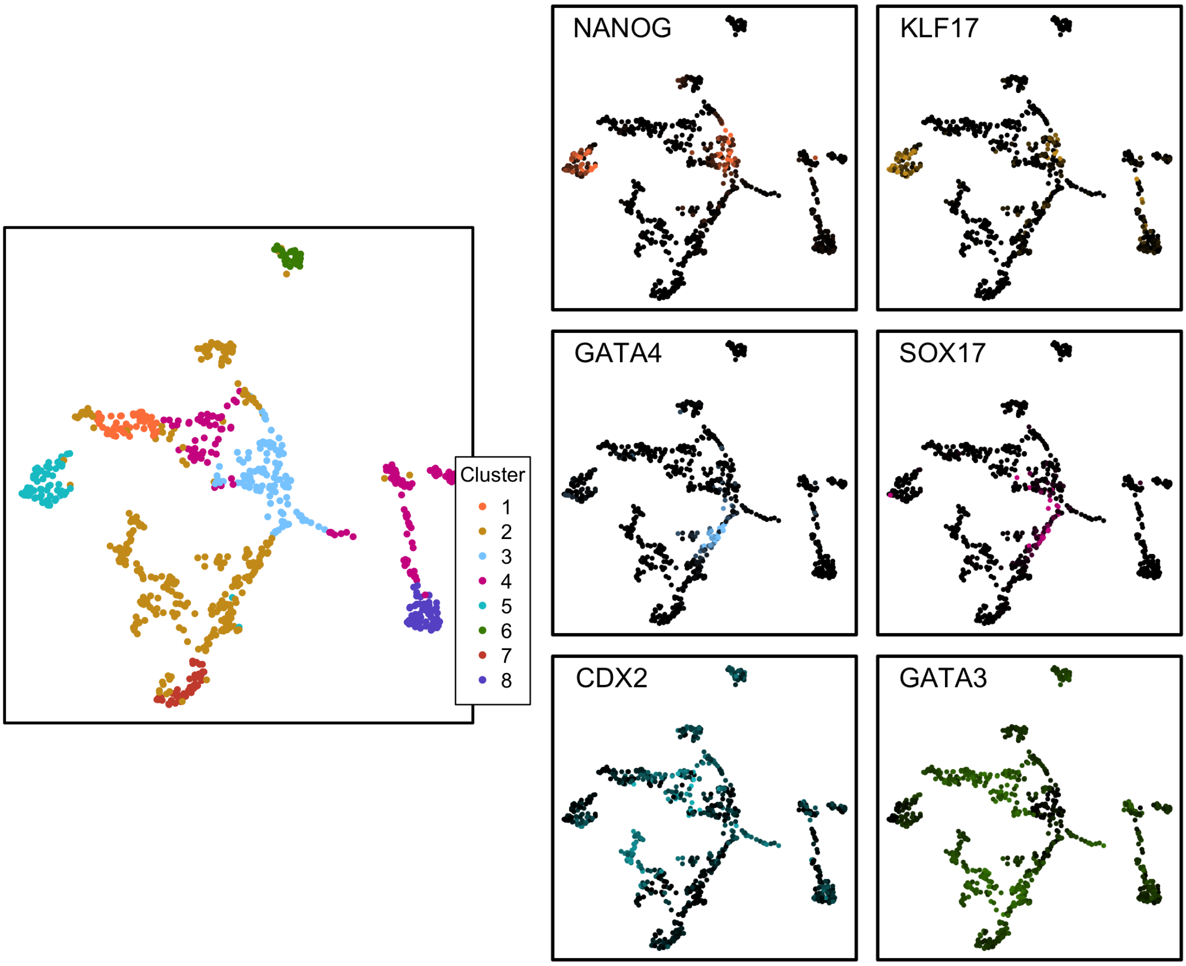


***Figure S1:*** *UMAP projections from the Laplacian eigenspace (UMAP-LE) [84, 86] show GMM-LE clusters and expression levels of validating marker genes for all cells from 5, 6, and 7 days post-fertilisation for the human embryonic development dataset [42].*

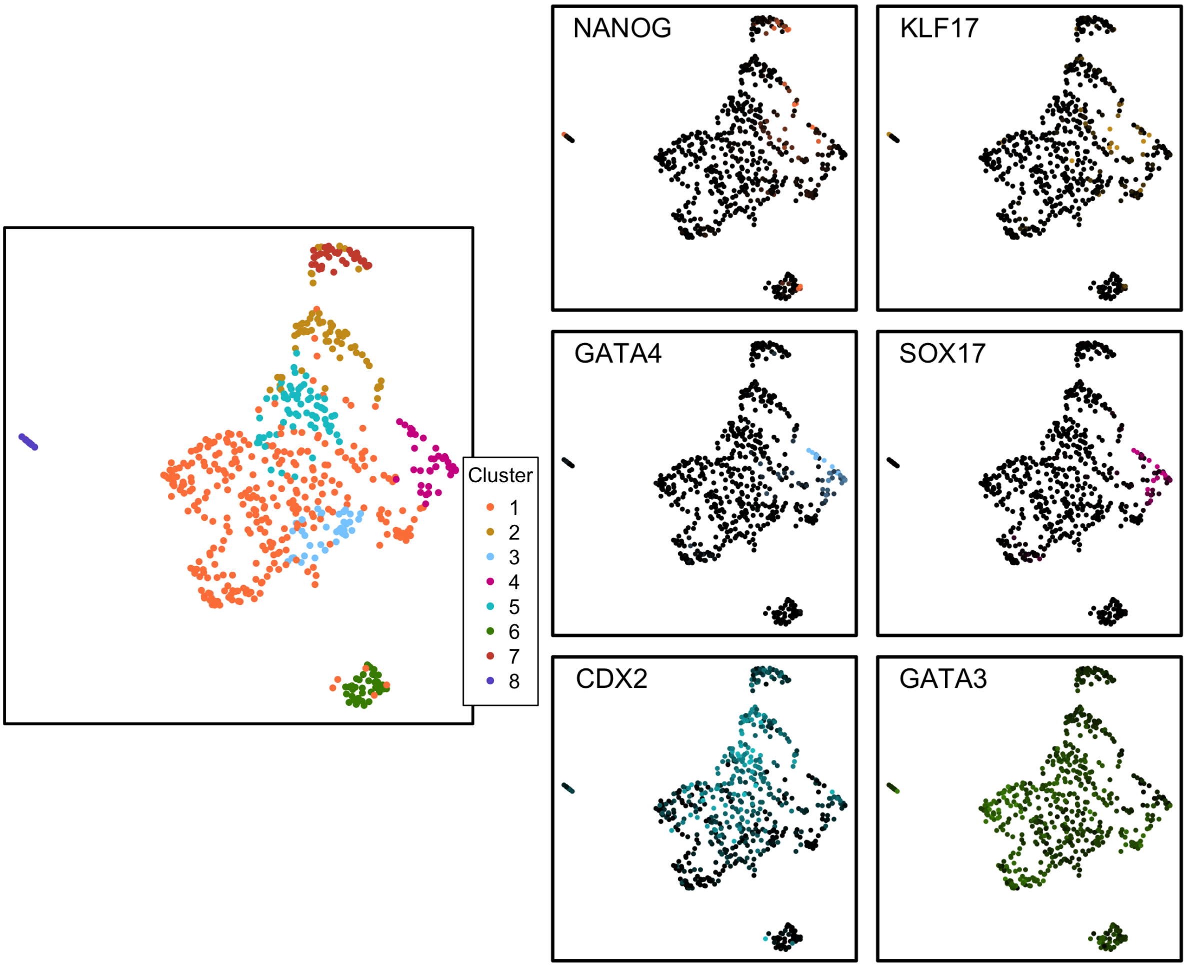


***Figure S2:*** *UMAP projections from the Laplacian eigenspace (UMAP-LE) [84, 86] show GMM-LE clusters and expression levels of validating marker genes, applied to cluster 2 from Figure S1, for the human embryonic development dataset [42].*

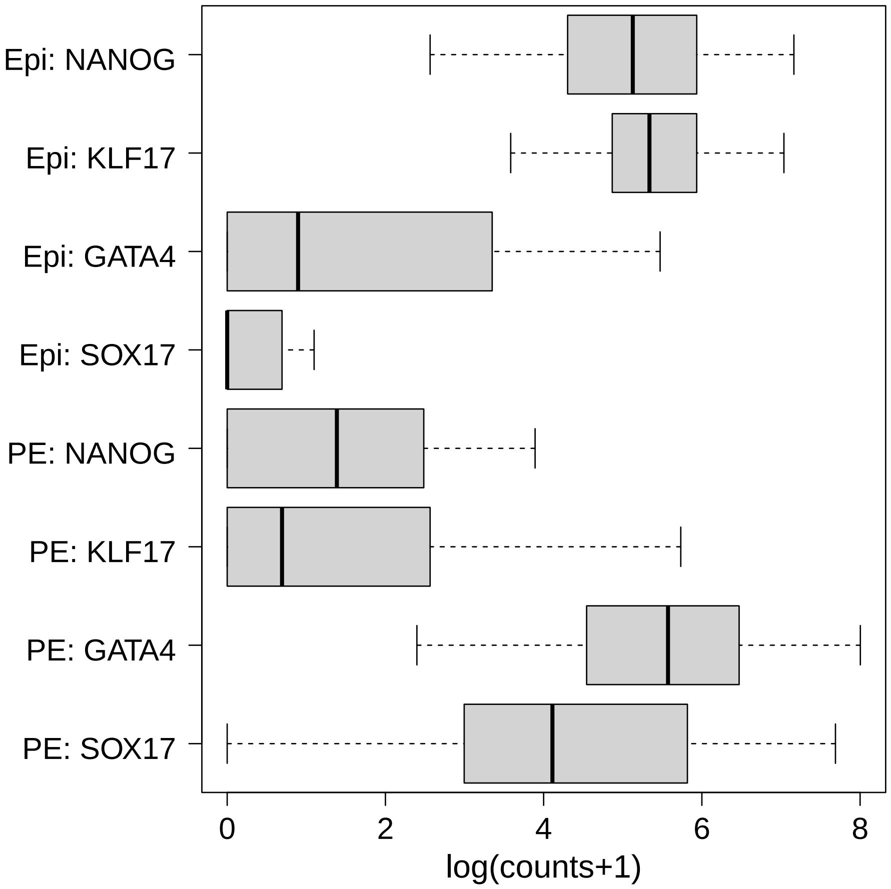


***Figure S3:*** *Boxplots showing marker genes for Epi (epiblast), validating the 68 Epi cells identified as cluster 5 in Figure S1, and for the PE (primitive endoderm), validating the 37 PE cells identified as cluster 4 in Figure S2, for the human embryonic development dataset [42].*

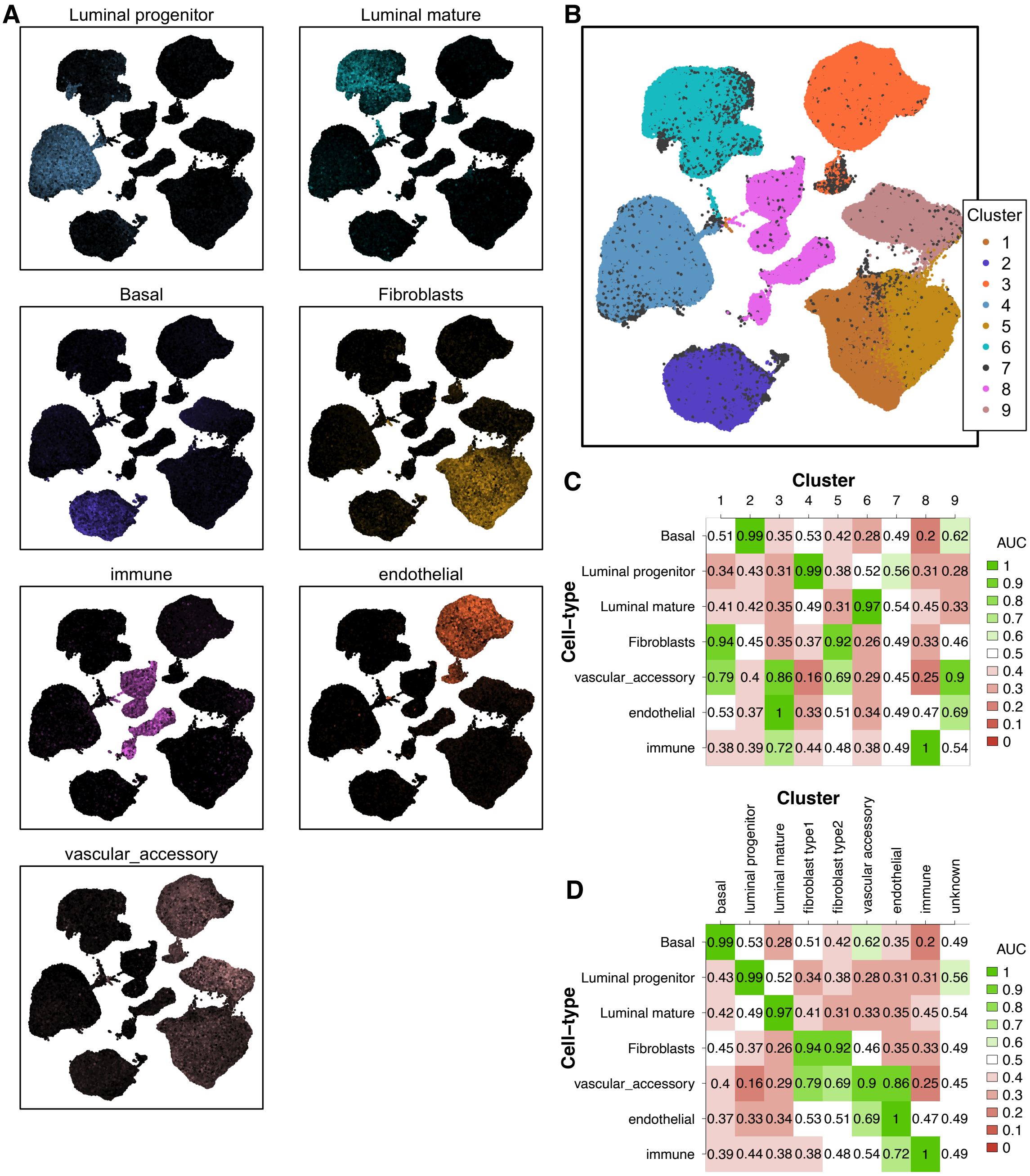


***Figure S4:*** *UMAP projections from the Laplacian eigenspace (UMAP-LE) [84,86] show (a) mean expression levels of validating marker genes and (b) GMM-LE clusters for all cells from the combined breast cancer at-risk datasets [52–58]. In (c) the AUC statistic is calculated (over cells) to compare membership of each cluster, with mean expression level of marker-genes for each cell-type. I.e., the AUC statistic quantifies how well the mean marker-gene expression for a cell-type predicts membership of a cluster. This allows automatic cell-type identification via maximum-AUC criterion as shown in (d), as well as assessment of clustering accuracy. These cell-type definitions are used subsequently in all analyses based on the combined breast cancer at-risk datasets.*

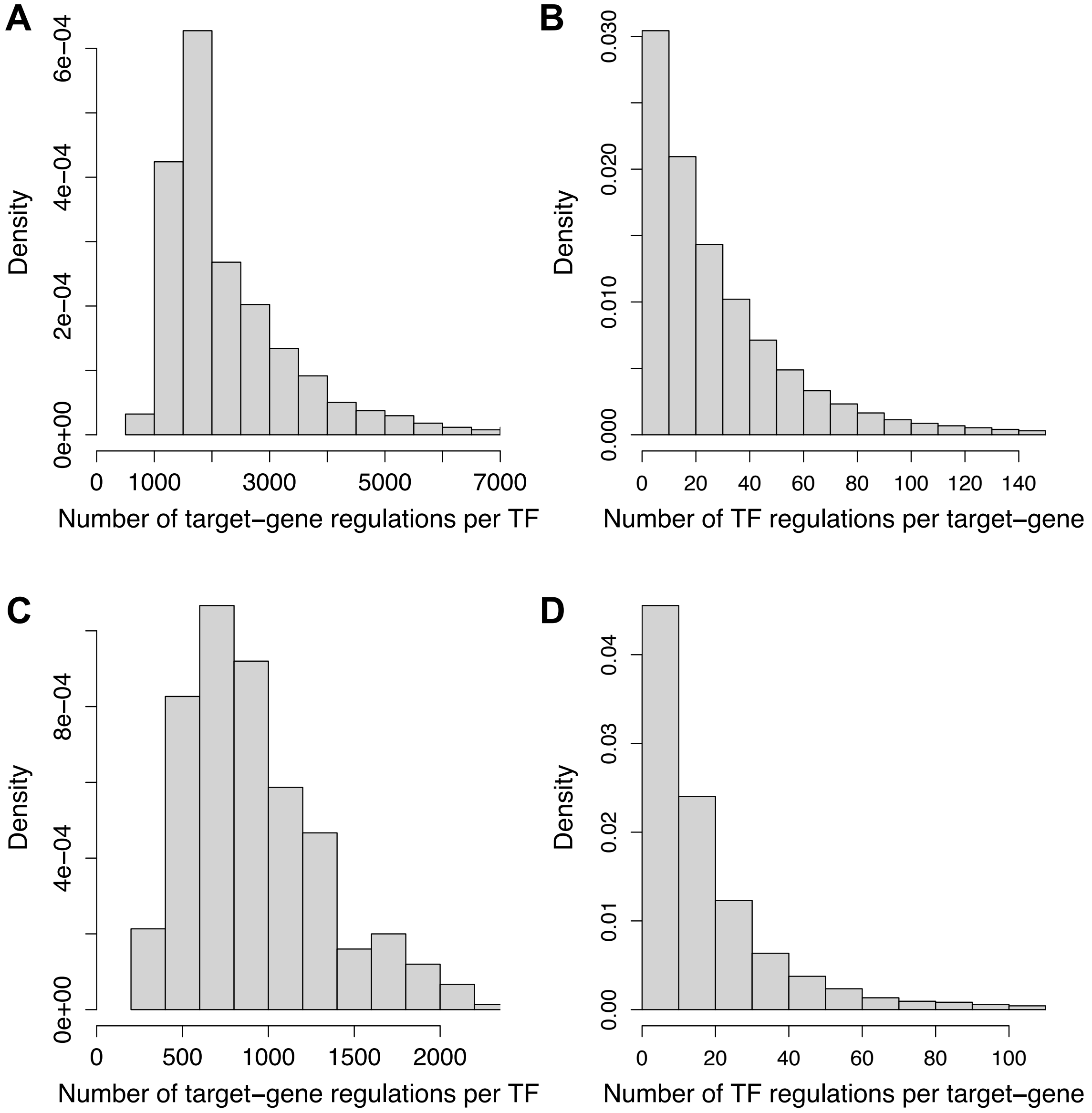


***Figure S5:*** *Histograms showing (a) and (c) the numbers of target-gene regulations for each TF, and (b) and (d) the number of TF regulators for each target-gene, in the epigenomic prior networks for (a) and (b) the breast cancer at-risk dataset, and (c) and (d) the human embryonic development dataset.*

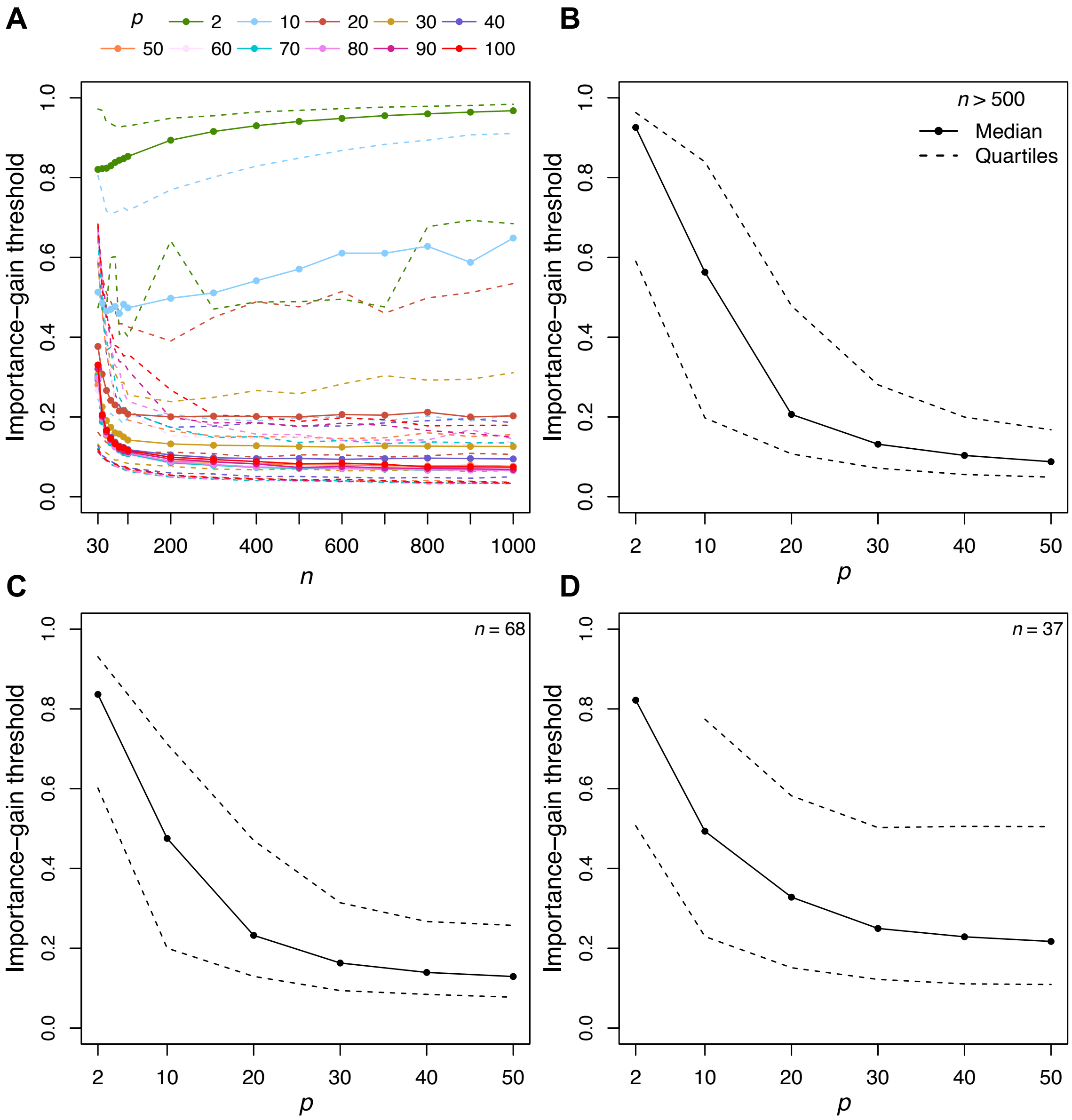


***Figure S6:*** *(a) Synthetic data with known ground-truth network structure are used to select the optimal importance-gain threshold (assessed by maximum-AUC criterion) for each target-gene in 250 synthetic datasets, for varying* $p$ *and* $n$*; solid lines show means, and dashed lines show lower and upper quartiles. For* $p > 50$ *there is little change in the optimal threshold, so we use the same threshold for all* $p > 50$*. Similarly for* $n > 500$ *(which includes all GRN inference based on the combined breast-cancer at-risk dataset), there is little change in the optimal threshold, so we use the same threshold for all* $n > 500$*, as shown in (b). For* $n < 500$*, we use thresholds specific to the sample size:* $n = 68$ *for epiblast, and* $n = 37$ *for primitive endoderm cells, as shown in (c) and (d) respectively.*

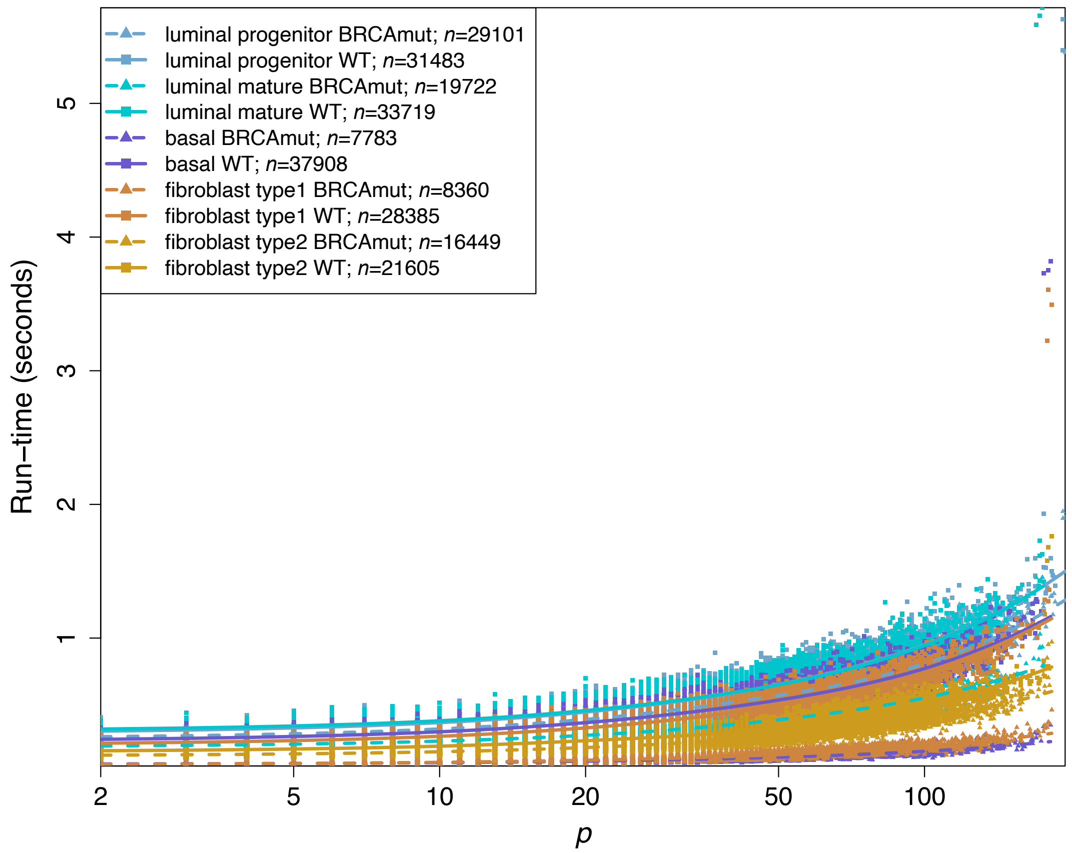


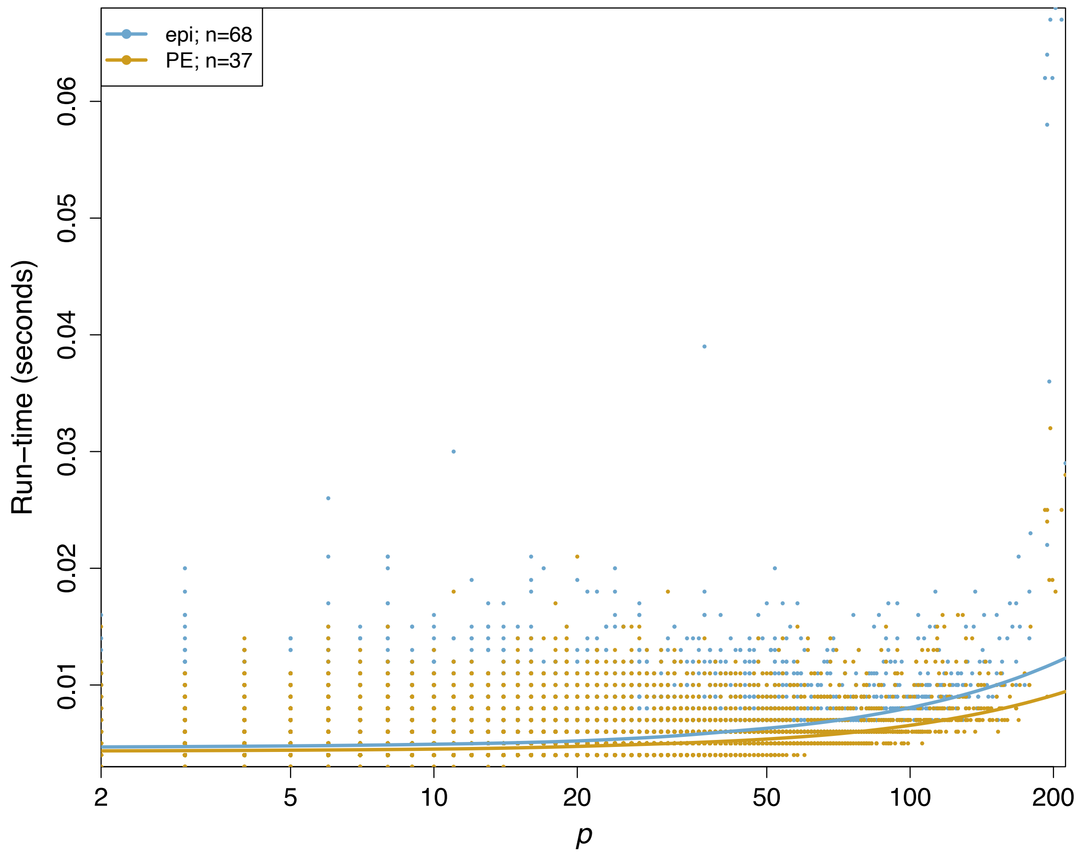


***Figure S7:*** *Runtime is plotted against number of TF predictors* p *in each target-gene random-forests model, for each GRN inference carried out for the combined breast-cancer at-risk dataset, and the human embryonic development dataset. Each point plotted corresponds to one random-forests model fit for one target-gene. Trend-lines are fitted with lowess regression.*

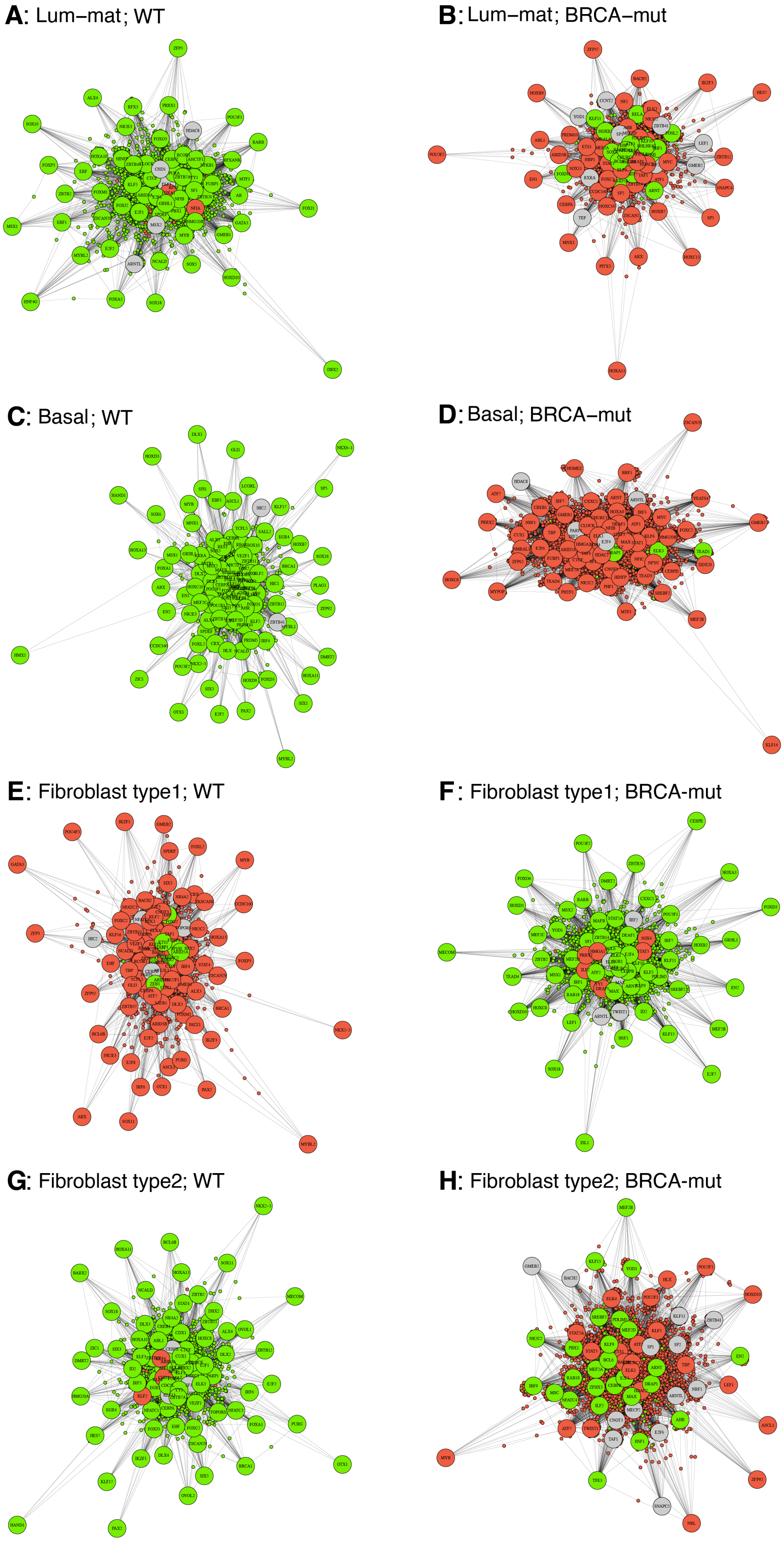


***Figure S8:*** *Differential regulation networks in (a) and (b) mature luminal cells (lum-mat), in (c) and (d) basal cells, and in (e), (f), (g), and (h) fibroblasts, for (a), (c), (e), and (g) BRCA1/2 wild-type (WT), and (b), (d), (f), and (h) BRCA1/2 mutation carriers (BRCAmut). Large circles indicate significantly differentially-regulating (diff-reg) transcription factors (Table S3, Table S4, Table S5, Table S6, Table S7); green and red indicate (respectively) significantly upregulated and downregulated TFs and genes; significance level is FDR* $p\text{-}val < 0.05$, *(t-test, Benjamini-Hochberg adjustment).*

| TF | OR | *p*-val | *p*-val RND |
|---|---|---|---|
| E4F1 | 1110 (851 - 1460) | <0.001 | <0.001 |
| BHLHE40 | 858 (644 - 1190) | <0.001 | <0.001 |
| CEBPZ | 844 (715 - 928) | <0.001 | <0.001 |
| NFYA | 738 (669 - 825) | <0.001 | <0.001 |
| NFYC | 733 (629 - 825) | <0.001 | <0.001 |
| ATF1 | 562 (456 - 716) | <0.001 | <0.001 |
| MITF | 482 (394 - 573) | <0.001 | <0.001 |
| ISX | 459 (316 - 629) | <0.001 | <0.001 |
| POU5F1 | 409 (311 - 530) | <0.001 | <0.001 |
| RFX2 | 394 (333 - 472) | <0.001 | <0.001 |
| CREM | 386 (309 - 475) | <0.001 | <0.001 |
| EN2 | 347 (231 - 505) | <0.001 | <0.001 |
| FOXI1 | 331 (286 - 372) | <0.001 | <0.001 |
| BARHL1 | 330 (172 - 588) | <0.001 | <0.001 |
| LHX6 | 296 (239 - 372) | <0.001 | <0.001 |
| TFE3 | 286 (237 - 338) | <0.001 | <0.001 |
| MAX | 257 (215 - 305) | <0.001 | <0.001 |
| ATF7 | 257 (199 - 330) | <0.001 | <0.001 |
| SOX9 | 256 (225 - 286) | <0.001 | <0.001 |
| KLF14 | 246 (230 - 262) | <0.001 | <0.001 |
| HMBOX1 | 243 (149 - 376) | <0.001 | <0.001 |
| ELK1 | 240 (214 - 274) | <0.001 | <0.001 |
| KLF11 | 233 (217 - 252) | <0.001 | <0.001 |
| GMEB2 | 230 (158 - 331) | <0.001 | <0.001 |
| CREB1 | 229 (191 - 272) | <0.001 | <0.001 |
| SP2 | 218 (207 - 233) | <0.001 | <0.001 |
| NRF1 | 212 (193 - 232) | <0.001 | <0.001 |
| E2F4 | 203 (191 - 218) | <0.001 | <0.001 |
| GSC | 200 (140 - 282) | <0.001 | <0.001 |
| ZBTB14 | 190 (172 - 211) | <0.001 | <0.001 |
| ATF2 | 184 (157 - 217) | <0.001 | <0.001 |
| ATF3 | 183 (153 - 215) | <0.001 | <0.001 |
| OVOL1 | 173 (98.6 - 287) | <0.001 | <0.001 |
| EGR1 | 173 (164 - 184) | <0.001 | <0.001 |
| YY1 | 166 (140 - 199) | <0.001 | <0.001 |
| PAX4 | 162 (101 - 250) | <0.001 | <0.001 |
| SNAI1 | 154 (118 - 201) | <0.001 | <0.001 |
| ERG | 154 (137 - 174) | <0.001 | <0.001 |
| ISL2 | 154 (108 - 217) | <0.001 | <0.001 |
| BARX1 | 147 (107 - 198) | <0.001 | <0.001 |
| E2F2 | 143 (121 - 167) | <0.001 | <0.001 |
| HOXA9 | 141 (62.4 - 286) | <0.001 | <0.001 |
| EMX1 | 139 (89.3 - 209) | <0.001 | <0.001 |
| ZBTB33 | 135 (104 - 175) | <0.001 | <0.001 |
| EGR4 | 127 (114 - 140) | <0.001 | <0.001 |
| E2F1 | 124 (114 - 135) | <0.001 | <0.001 |
| ETV6 | 123 (109 - 139) | <0.001 | <0.001 |
| ELF1 | 116 (105 - 128) | <0.001 | <0.001 |
| HBP1 | 115 (70.5 - 178) | <0.001 | <0.001 |
| SPDEF | 112 (90.6 - 137) | <0.001 | <0.001 |
| SOX13 | 111 (65.1 - 180) | <0.001 | <0.001 |
| JUN | 111 (89.8 - 137) | <0.001 | <0.001 |
| ELK3 | 111 (91.1 - 134) | <0.001 | <0.001 |
| HOXD13 | 110 (77.9 - 153) | <0.001 | <0.001 |
| EHF | 110 (97.6 - 124) | <0.001 | <0.001 |
| AHR | 109 (65.9 - 172) | <0.001 | <0.001 |
| ERF | 107 (76.5 - 147) | <0.001 | <0.001 |
| SREBF2 | 102 (75.7 - 135) | <0.001 | <0.001 |
| HOXA13 | 102 (70.2 - 144) | <0.001 | <0.001 |
| PRRX2 | 100 (72.1 - 137) | <0.001 | <0.001 |
| BSX | 97 (69 - 134) | <0.001 | <0.001 |
| EVX1 | 93.9 (46.1 - 172) | <0.001 | <0.001 |
| TAF1 | 91.2 (83.3 - 99.8) | <0.001 | <0.001 |
| CDX1 | 88.5 (61.7 - 124) | <0.001 | <0.001 |
| FOXN3 | 87.2 (51.6 - 139) | <0.001 | <0.001 |
| LHX2 | 86.8 (65.6 - 113) | <0.001 | <0.001 |
| KLF16 | 83.3 (73.7 - 93.4) | <0.001 | <0.001 |
| PRRX1 | 83.2 (65.1 - 105) | <0.001 | <0.001 |
| ETS1 | 83 (69.4 - 98.5) | <0.001 | <0.001 |
| CDC5L | 82.2 (34.5 - 169) | <0.001 | <0.001 |
| MEF2D | 81.9 (52.5 - 122) | <0.001 | <0.001 |
| ARNT | 80.2 (51.4 - 120) | <0.001 | <0.001 |
| PBX1 | 79.7 (61.3 - 102) | <0.001 | <0.001 |
| NKX2-3 | 78 (43.9 - 129) | <0.001 | <0.001 |
| WT1 | 77.6 (70.5 - 85.6) | <0.001 | <0.001 |
| NKX6-2 | 77.3 (48 - 119) | <0.001 | <0.001 |
| LHX8 | 77 (46.3 - 122) | <0.001 | <0.001 |
| LHX3 | 75 (39.1 - 132) | <0.001 | <0.001 |
| BARX2 | 74 (45.7 - 114) | <0.001 | <0.001 |
| TBP | 72.6 (46.5 - 109) | <0.001 | <0.001 |
| TEF | 71.3 (40.9 - 116) | <0.001 | <0.001 |
| IRF2 | 67.1 (52.6 - 85) | <0.001 | <0.001 |
| SOX5 | 66.9 (44.6 - 97.2) | <0.001 | <0.001 |
| MEF2C | 66.5 (41.1 - 103) | <0.001 | <0.001 |
| CEBPB | 65.9 (42.1 - 99.1) | <0.001 | <0.001 |
| FOXO1 | 63.7 (39.5 - 97.9) | <0.001 | <0.001 |
| GATA1 | 63 (46.4 - 83.9) | <0.001 | <0.001 |
| ARX | 62.5 (36.2 - 102) | <0.001 | <0.001 |
| CUX1 | 62 (36.4 - 100) | <0.001 | <0.001 |
| ZSCAN29 | 61.6 (34.3 - 103) | <0.001 | <0.001 |
| MSX2 | 60.9 (39.1 - 91.1) | <0.001 | <0.001 |
| ZFHX3 | 57.9 (26.1 - 112) | <0.001 | <0.001 |
| LMX1A | 57.7 (33.2 - 93.8) | <0.001 | <0.001 |
| POU4F3 | 56.4 (29.5 - 99.1) | <0.001 | <0.001 |
| FOXJ3 | 55 (41.1 - 72.4) | <0.001 | <0.001 |
| CEBPA | 52.8 (36.1 - 75) | <0.001 | <0.001 |
| CDX2 | 52.5 (38.9 - 69.6) | <0.001 | <0.001 |
| IRF7 | 52.4 (38.5 - 69.7) | <0.001 | <0.001 |
| ALX3 | 52 (38.3 - 69.6) | <0.001 | <0.001 |
| KLF8 | 51.8 (39.6 - 66.9) | <0.001 | <0.001 |
| STAT1 | 51.2 (41.6 - 62.6) | <0.001 | <0.001 |
| CLOCK | 50.7 (36.5 - 69.1) | <0.001 | <0.001 |
| HOXA10 | 50.6 (31 - 78.8) | <0.001 | <0.001 |
| ONECUT2 | 49.7 (21 - 101) | <0.001 | <0.001 |
| FOXG1 | 48.8 (36 - 64.9) | <0.001 | <0.001 |
| ZBTB26 | 48.2 (34.2 - 66.4) | <0.001 | <0.001 |
| SREBF1 | 48.2 (29.1 - 75.1) | <0.001 | <0.001 |
| FOSL2 | 47.8 (29.9 - 72.4) | <0.001 | <0.001 |
| POU2F1 | 47 (34.9 - 62.3) | <0.001 | <0.001 |
| ELF3 | 46.3 (39.9 - 53.8) | <0.001 | <0.001 |
| MZF1 | 46.1 (38.7 - 54.9) | <0.001 | <0.001 |
| RXRA | 45.5 (38.1 - 54.1) | <0.001 | <0.001 |
| HOXD10 | 45.2 (16.2 - 102) | <0.001 | <0.001 |
| HOXC10 | 44.8 (16.1 - 101) | <0.001 | <0.001 |
| DLX3 | 44.1 (27.1 - 68.6) | <0.001 | <0.001 |
| OTX1 | 43.8 (27.7 - 66) | <0.001 | <0.001 |
| OLIG3 | 42.8 (18.2 - 86.3) | <0.001 | <0.001 |
| DLX1 | 42.2 (13.6 - 99.9) | <0.001 | <0.001 |
| CRX | 41.7 (27.3 - 61.5) | <0.001 | <0.001 |
| SOX14 | 41.7 (16.6 - 87.4) | <0.001 | <0.001 |
| REST | 41.6 (23.5 - 68.5) | <0.001 | <0.001 |
| BPTF | 38.9 (26.9 - 54.3) | <0.001 | <0.001 |
| GLI2 | 37.6 (28 - 49.5) | <0.001 | <0.001 |
| HOXD3 | 37.3 (20.1 - 63.7) | <0.001 | <0.001 |
| NR1H4 | 37 (27.7 - 48.5) | <0.001 | <0.001 |
| VSX2 | 36.9 (20.8 - 60.8) | <0.001 | <0.001 |
| EN1 | 36.6 (20.1 - 61.6) | <0.001 | <0.001 |
| GBX2 | 36.5 (20.6 - 60.3) | <0.001 | <0.001 |
| FOXA1 | 36.5 (27.3 - 48) | <0.001 | <0.001 |
| BRCA1 | 34.6 (23.5 - 49.3) | <0.001 | <0.001 |
| BCL6 | 34.3 (23 - 49.3) | <0.001 | <0.001 |
| ALX1 | 33.8 (15.2 - 65.5) | <0.001 | <0.001 |
| FOXD2 | 33.5 (19 - 55.1) | <0.001 | <0.001 |
| ONECUT3 | 33.5 (4.01 - 123) | 0.00176 | <0.001 |
| PAX3 | 33.1 (14.1 - 67) | <0.001 | <0.001 |
| ZBTB12 | 33 (18.9 - 53.8) | <0.001 | <0.001 |
| RARG | 32.5 (25.3 - 41.3) | <0.001 | <0.001 |
| SOX10 | 31.5 (25.3 - 38.7) | <0.001 | <0.001 |
| LHX1 | 29.3 (13.8 - 55) | <0.001 | <0.001 |
| FOXC2 | 26.8 (17.8 - 38.8) | <0.001 | <0.001 |
| FOXM1 | 26.6 (15.6 - 42.4) | <0.001 | <0.001 |
| LEF1 | 26.3 (19.2 - 35.2) | <0.001 | <0.001 |
| POU3F2 | 25.8 (16.8 - 37.9) | <0.001 | <0.001 |
| FOXC1 | 25.5 (18.1 - 34.8) | <0.001 | <0.001 |
| SIX1 | 25.4 (9.25 - 55.9) | <0.001 | <0.001 |
| TLX2 | 24.8 (6.7 - 64.6) | <0.001 | <0.001 |
| POU3F1 | 24.7 (16.3 - 36.2) | <0.001 | <0.001 |
| SOX7 | 24.5 (2.94 - 90.6) | 0.00321 | <0.001 |
| DLX2 | 24.2 (10.2 - 49) | <0.001 | <0.001 |
| HOXB7 | 23.9 (6.44 - 62.4) | <0.001 | <0.001 |
| SCRT1 | 23.6 (13.4 - 38.7) | <0.001 | <0.001 |
| IRF4 | 23.1 (11.5 - 41.8) | <0.001 | <0.001 |
| RARB | 22.5 (14.6 - 33.4) | <0.001 | <0.001 |
| ALX4 | 22.2 (8.04 - 49.2) | <0.001 | <0.001 |
| FOXL1 | 22 (12.7 - 35.6) | <0.001 | <0.001 |
| PROP1 | 21.4 (8.53 - 44.7) | <0.001 | <0.001 |
| NR2E1 | 20.6 (1 - 117) | 0.0476 | 0.012 |
| STAT3 | 19.6 (4.01 - 57.9) | <0.001 | <0.001 |
| SOX1 | 14.9 (4.79 - 35.5) | <0.001 | <0.001 |
| NR2E3 | 13.5 (6.14 - 25.9) | <0.001 | <0.001 |
| FOXJ2 | 13.4 (6.39 - 24.8) | <0.001 | <0.001 |
| HESX1 | 13.4 (3.62 - 34.6) | <0.001 | <0.001 |
| NFE2L1 | 13.3 (7 - 23) | <0.001 | <0.001 |
| HMGA1 | 13.2 (4.25 - 31.5) | <0.001 | <0.001 |
| GRHL1 | 12 (5.44 - 22.9) | <0.001 | <0.001 |
| PHOX2B | 11.9 (1.43 - 43.7) | 0.0128 | <0.001 |
| DLX4 | 11 (2.25 - 32.8) | 0.00282 | <0.001 |
| NANOG | 9.57 (3.83 - 19.8) | <0.001 | <0.001 |
| MYBL2 | 1 (1 - 135) | 1 | 1 |
| HOXB9 | 1 (1 - 48.9) | 1 | 1 |

***Table S1:*** *Odds-ratios (with 95% C.I.s) as displayed in Figure 4 resulting from comparison of the overlap between the DNA regions inferred as regulated by 170 TFs in the inner cell mass (ICM) according to both the modified-lever-cistarget (MLC) method and the RGT-HINT [36] method, together with significance p-vals calculated according to Fisher's exact test, as well as p-vals calculated by randomising the chromosome data and comparing the resulting resampled odd-ratios with the observed odds-ratios.*

**A**

| TF | mean diff-reg stat | *p*-val | adj *p*-val |
|---|---|---|---|
| E2F1 | 0.00756 | <0.001 | <0.001 |
| BCLAF1 | 0.0045 | <0.001 | <0.001 |
| NANOG | 0.0043 | <0.001 | <0.001 |
| ETS1 | 0.00292 | <0.001 | <0.001 |
| ARX | 0.00265 | <0.001 | <0.001 |
| IRF4 | 0.00245 | <0.001 | <0.001 |
| KLF17 | 0.00198 | <0.001 | <0.001 |
| BRCA1 | 0.00172 | <0.001 | <0.001 |
| KLF11 | 0.00162 | <0.001 | <0.001 |
| LHX8 | 0.00159 | <0.001 | <0.001 |
| KLF4 | 0.00159 | <0.001 | <0.001 |
| DLX3 | 0.00159 | 0.001 | 0.005 |
| MZF1 | 0.00159 | <0.001 | <0.001 |
| ARID5A | 0.00152 | <0.001 | <0.001 |
| FOXC1 | 0.00145 | <0.001 | 0.003 |
| GLI2 | 0.00129 | <0.001 | <0.001 |
| ALX1 | 0.00124 | 0.017 | 0.047 |
| BBX | 0.00123 | <0.001 | 0.002 |
| CNOT3 | 0.00114 | <0.001 | 0.002 |
| KLF2 | 0.00112 | <0.001 | <0.001 |
| HMGA1 | 0.0011 | 0.011 | 0.034 |
| SF1 | 0.00108 | <0.001 | 0.003 |
| PBX1 | 0.00107 | <0.001 | <0.001 |
| GRHL1 | 0.00107 | <0.001 | 0.002 |
| HDAC2 | 0.00104 | 0.008 | 0.027 |
| NRF1 | 0.000995 | 0.004 | 0.014 |
| CXXC1 | 0.000949 | 0.002 | 0.007 |
| MEF2D | 0.000906 | <0.001 | <0.001 |
| POU3F1 | 0.000869 | <0.001 | <0.001 |
| DLX4 | 0.000848 | <0.001 | 0.002 |
| CDX1 | 0.000838 | 0.012 | 0.035 |
| SOX1 | 0.000795 | <0.001 | <0.001 |
| GATA2 | 0.000776 | 0.004 | 0.014 |
| BRF1 | 0.000761 | 0.014 | 0.04 |
| SOX15 | 0.000759 | <0.001 | <0.001 |
| GBX2 | 0.000759 | <0.001 | 0.002 |
| PATZ1 | 0.000741 | 0.001 | 0.006 |
| YBX1 | 0.000736 | 0.012 | 0.035 |
| FOXN3 | 0.000734 | <0.001 | 0.002 |
| FOXG1 | 0.000713 | <0.001 | <0.001 |
| KLF1 | 0.000711 | <0.001 | <0.001 |
| ELF3 | 0.000693 | 0.007 | 0.023 |
| DMBX1 | 0.000667 | <0.001 | <0.001 |
| MBD2 | 0.000653 | 0.016 | 0.045 |
| HESX1 | 0.000632 | 0.002 | 0.009 |
| PURA | 0.000631 | 0.011 | 0.033 |
| PARP1 | 0.000631 | 0.011 | 0.034 |
| RCOR1 | 0.000628 | 0.011 | 0.034 |
| ONECUT2 | 0.000609 | 0.003 | 0.012 |
| SCRT1 | 0.000586 | <0.001 | <0.001 |
| MYB | 0.000586 | <0.001 | <0.001 |
| ATF7 | 0.000548 | 0.007 | 0.023 |
| ZFP1 | 0.000539 | 0.007 | 0.023 |
| TBPL2 | 0.000492 | 0.006 | 0.021 |
| POU6F1 | 0.000442 | 0.001 | 0.005 |
| BARX1 | 0.000413 | <0.001 | <0.001 |
| BARX2 | 0.000399 | 0.006 | 0.02 |
| AHDC1 | 0.000385 | 0.004 | 0.014 |
| SIX3 | 0.000347 | <0.001 | 0.002 |
| FOXL2 | 0.000344 | <0.001 | 0.004 |
| OLIG3 | 0.000337 | <0.001 | <0.001 |
| KLF14 | 0.00032 | <0.001 | <0.001 |
| EMX1 | 0.000298 | 0.007 | 0.024 |
| PRRX1 | 0.000281 | <0.001 | <0.001 |
| HOXA13 | 0.00028 | <0.001 | <0.001 |
| TLX2 | 0.000145 | <0.001 | <0.001 |
| MYOD1 | 0.000138 | <0.001 | 0.002 |
| POU4F2 | 7.2e-05 | 0.009 | 0.029 |

**B**

| TF | mean diff-reg stat | *p*-val | adj *p*-val |
|---|---|---|---|
| ARID3A | 0.0134 | <0.001 | <0.001 |
| EGR1 | 0.0112 | <0.001 | <0.001 |
| ELF1 | 0.0069 | <0.001 | <0.001 |
| PRDM11 | 0.00256 | <0.001 | <0.001 |
| E2F2 | 0.00233 | <0.001 | 0.002 |
| ATF3 | 0.00174 | <0.001 | <0.001 |
| CRX | 0.00172 | <0.001 | <0.001 |
| SPDEF | 0.00163 | <0.001 | <0.001 |
| HLX | 0.0014 | <0.001 | <0.001 |
| EHF | 0.00138 | <0.001 | <0.001 |
| NR2E3 | 0.00132 | <0.001 | <0.001 |
| LIN28B | 0.00121 | 0.002 | 0.009 |
| EN1 | 0.0012 | <0.001 | 0.003 |
| HOXA9 | 0.00118 | <0.001 | <0.001 |
| CREB1 | 0.00105 | <0.001 | 0.003 |
| NFIB | 0.00101 | <0.001 | <0.001 |
| ZFP62 | 0.000916 | 0.017 | 0.048 |
| HOXA3 | 0.000913 | <0.001 | <0.001 |
| ZKSCAN7 | 0.000906 | <0.001 | <0.001 |
| HOXA10 | 0.000885 | <0.001 | 0.003 |
| GSC | 0.00086 | <0.001 | <0.001 |
| RARA | 0.000808 | <0.001 | 0.004 |
| NR1H4 | 0.000797 | <0.001 | <0.001 |
| JUN | 0.000744 | <0.001 | 0.002 |
| VSX2 | 0.000596 | 0.002 | 0.007 |
| SOX14 | 0.000596 | <0.001 | 0.003 |
| PITX3 | 0.00056 | <0.001 | <0.001 |
| SIX1 | 0.000557 | 0.012 | 0.036 |
| RARB | 0.000507 | 0.003 | 0.011 |
| NKX2-5 | 0.000487 | 0.002 | 0.007 |
| OTP | 0.000481 | 0.005 | 0.016 |
| EN2 | 0.000279 | <0.001 | 0.002 |
| MEF2B | 0.000251 | 0.015 | 0.043 |
| ISX | 0.000158 | 0.008 | 0.026 |
| HOXC8 | 0.000123 | <0.001 | <0.001 |

***Table S2:*** *Transcription factors with significant differential regulation (FDR $p\text{-}val < 0.05$, t-test, Benjamini-Hochberg adjustment) showing increased regulation (hyper-regulation) in (a) epiblast and (b) primitive endoderm cells.*

**A**

| TF | mean diff-reg stat | *p*-val | adj *p*-val |
|---|---:|---:|---:|
| E2F1 | 0.00783 | <0.001 | <0.001 |
| ELK1 | 0.00577 | <0.001 | <0.001 |
| E2F4 | 0.00519 | <0.001 | <0.001 |
| CEBPB | 0.00359 | <0.001 | <0.001 |
| SOX10 | 0.00358 | <0.001 | <0.001 |
| BPTF | 0.00322 | <0.001 | <0.001 |
| EHF | 0.0021 | <0.001 | <0.001 |
| ZBTB7B | 0.00203 | <0.001 | <0.001 |
| ARNTL | 0.00201 | <0.001 | <0.001 |
| HINFP | 0.00178 | <0.001 | <0.001 |
| FOXJ2 | 0.00168 | <0.001 | <0.001 |
| PBX1 | 0.00152 | <0.001 | <0.001 |
| NFIB | 0.00142 | <0.001 | <0.001 |
| ZBTB20 | 0.00142 | <0.001 | <0.001 |
| NFATC1 | 0.00137 | <0.001 | <0.001 |
| FOXP4 | 0.00134 | <0.001 | <0.001 |
| YY1 | 0.00132 | <0.001 | <0.001 |
| CLOCK | 0.00132 | <0.001 | <0.001 |
| GRHL1 | 0.00125 | <0.001 | 0.001 |
| PURA | 0.00125 | <0.001 | <0.001 |
| DLX2 | 0.00114 | <0.001 | <0.001 |
| RCOR1 | 0.00113 | <0.001 | <0.001 |
| AHDC1 | 0.00111 | <0.001 | <0.001 |
| SF1 | 0.00111 | <0.001 | <0.001 |
| SATB1 | 0.00109 | <0.001 | <0.001 |
| ETV4 | 0.00103 | <0.001 | <0.001 |
| TFCP2L1 | 0.00102 | <0.001 | <0.001 |
| NFIA | 0.000999 | <0.001 | <0.001 |
| ZKSCAN8 | 0.000982 | <0.001 | <0.001 |
| MTF1 | 0.000889 | <0.001 | <0.001 |
| NCALD | 0.000859 | <0.001 | <0.001 |
| CHD1 | 0.000782 | <0.001 | <0.001 |
| ERF | 0.000759 | <0.001 | <0.001 |
| SPDEF | 0.000757 | <0.001 | 0.003 |
| ZBTB7A | 0.000747 | <0.001 | <0.001 |
| HMG20B | 0.000744 | <0.001 | <0.001 |
| MYBL2 | 0.000718 | <0.001 | <0.001 |
| FOXM1 | 0.000703 | <0.001 | <0.001 |
| SOX5 | 0.000694 | <0.001 | <0.001 |
| E2F2 | 0.000692 | <0.001 | <0.001 |
| TAF6 | 0.000681 | <0.001 | <0.001 |
| ZFP62 | 0.000664 | 0.005 | 0.012 |
| IRF4 | 0.000658 | <0.001 | <0.001 |
| NR2C2 | 0.000658 | 0.005 | 0.013 |
| CXXC1 | 0.000632 | <0.001 | <0.001 |
| E2F6 | 0.000598 | <0.001 | 0.003 |
| LCORL | 0.000567 | <0.001 | <0.001 |
| FOXA3 | 0.000536 | <0.001 | <0.001 |
| PRDM4 | 0.000534 | <0.001 | <0.001 |
| RFX3 | 0.000497 | 0.002 | 0.005 |
| NFIC | 0.000496 | 0.013 | 0.029 |
| ATF2 | 0.000411 | 0.005 | 0.012 |
| IRF3 | 0.000405 | 0.003 | 0.007 |
| DLX3 | 0.000396 | 0.012 | 0.027 |
| HES7 | 0.000369 | <0.001 | <0.001 |
| FUBP1 | 0.000358 | 0.009 | 0.02 |
| SALL2 | 0.000329 | 0.001 | 0.004 |
| PHTF1 | 0.000314 | 0.009 | 0.021 |
| AHCTF1 | 0.000313 | 0.011 | 0.024 |
| HDAC8 | 0.000309 | 0.005 | 0.012 |
| HMG20A | 0.000294 | 0.006 | 0.014 |
| ERG | 0.000268 | 0.002 | 0.004 |
| TEF | 0.000265 | 0.002 | 0.005 |
| ZKSCAN3 | 0.000259 | <0.001 | <0.001 |
| HOXC8 | 0.000253 | 0.012 | 0.027 |
| MYB | 0.000248 | 0.002 | 0.006 |
| SMAD6 | 0.000227 | <0.001 | 0.002 |
| FOXJ1 | 0.000193 | <0.001 | <0.001 |
| FOXL2 | 0.000179 | <0.001 | 0.002 |
| SPIB | 0.000171 | <0.001 | <0.001 |
| PAX2 | 0.000141 | <0.001 | <0.001 |
| ZFP92 | 0.000139 | <0.001 | <0.001 |
| FOXP3 | 0.000129 | 0.003 | 0.008 |
| HAND1 | 0.000128 | 0.004 | 0.01 |
| KLF14 | 0.000106 | 0.017 | 0.036 |

**B**

| TF | mean diff-reg stat | *p*-val | adj *p*-val |
|---|---:|---:|---:|
| BACH1 | 0.00373 | <0.001 | <0.001 |
| BCLAF1 | 0.00363 | <0.001 | <0.001 |
| FOXC2 | 0.00281 | <0.001 | <0.001 |
| MAX | 0.00278 | <0.001 | <0.001 |
| ATF3 | 0.00278 | <0.001 | <0.001 |
| ELF1 | 0.00277 | <0.001 | <0.001 |
| HDAC2 | 0.00262 | <0.001 | <0.001 |
| STAT1 | 0.00261 | <0.001 | <0.001 |
| FOXC1 | 0.00259 | <0.001 | <0.001 |
| ETS1 | 0.00246 | <0.001 | <0.001 |
| SP100 | 0.00235 | <0.001 | <0.001 |
| CHURC1 | 0.00226 | <0.001 | <0.001 |
| ELK3 | 0.00212 | <0.001 | <0.001 |
| NFKB1 | 0.00201 | <0.001 | <0.001 |
| ELF3 | 0.00186 | <0.001 | <0.001 |
| PRRX2 | 0.00183 | <0.001 | <0.001 |
| KLF10 | 0.00151 | <0.001 | <0.001 |
| ARID3A | 0.00146 | 0.001 | 0.003 |
| TAF1 | 0.00143 | <0.001 | <0.001 |
| IRF6 | 0.00141 | <0.001 | <0.001 |
| SOX4 | 0.00136 | <0.001 | <0.001 |
| BCL6 | 0.00132 | <0.001 | <0.001 |
| SP1 | 0.00122 | <0.001 | <0.001 |
| RELA | 0.00109 | <0.001 | <0.001 |
| HMGA1 | 0.00109 | <0.001 | 0.001 |
| ID2 | 0.00109 | <0.001 | <0.001 |
| IRF2 | 0.001 | <0.001 | <0.001 |
| CEBPA | 0.000999 | <0.001 | <0.001 |
| PDLIM5 | 0.000987 | <0.001 | <0.001 |
| MEF2A | 0.000924 | <0.001 | <0.001 |
| MTA3 | 0.000789 | <0.001 | <0.001 |
| ZFP57 | 0.000773 | <0.001 | <0.001 |
| YOD1 | 0.000753 | <0.001 | <0.001 |
| ARID5A | 0.000745 | 0.003 | 0.009 |
| IRF1 | 0.000726 | 0.003 | 0.007 |
| RARB | 0.000724 | <0.001 | <0.001 |
| PHF1 | 0.000684 | <0.001 | <0.001 |
| ELK4 | 0.000639 | <0.001 | <0.001 |
| ZSCAN2 | 0.000593 | <0.001 | <0.001 |
| MLX | 0.000567 | <0.001 | <0.001 |
| HBP1 | 0.000552 | <0.001 | <0.001 |
| VEZF1 | 0.000537 | <0.001 | 0.002 |
| MSX2 | 0.000486 | <0.001 | <0.001 |
| KLF13 | 0.000484 | 0.013 | 0.027 |
| SP2 | 0.000474 | <0.001 | <0.001 |
| HSF1 | 0.000437 | 0.005 | 0.013 |
| HOXB2 | 0.000427 | <0.001 | <0.001 |
| IRF9 | 0.000421 | 0.004 | 0.01 |
| ARID5B | 0.000405 | 0.008 | 0.019 |
| MYBL1 | 0.000392 | 0.013 | 0.028 |
| CNOT3 | 0.000358 | 0.018 | 0.038 |
| MYC | 0.000348 | 0.005 | 0.013 |
| RXRA | 0.000347 | 0.01 | 0.022 |
| FOXJ3 | 0.000346 | <0.001 | 0.002 |
| MEF2B | 0.000344 | <0.001 | <0.001 |
| STAT4 | 0.000306 | 0.002 | 0.005 |
| FOXO3 | 0.000292 | 0.019 | 0.04 |
| ZFHX3 | 0.000255 | 0.022 | 0.047 |
| NKX2-5 | 0.000207 | 0.007 | 0.016 |
| ZIC1 | 0.000187 | <0.001 | <0.001 |
| FOXD3 | 0.00015 | <0.001 | <0.001 |
| CDX1 | 0.000148 | 0.004 | 0.011 |

***Table S3:*** *Transcription factors with significant differential regulation (FDR* $p\text{-}val < 0.05$*, t-test, Benjamini-Hochberg adjustment) in luminal progenitor cells that are (a) hypo-regulating and (b) hyper-regulating in BRCA1/2 mutation carriers (BRCAmut) compared to BRCA1/2 wild-type (WT).*

**A**

| TF | mean diff-reg stat | *p*-val | adj *p*-val |
|---|---:|---:|---:|
| BPTF | 0.0133 | <0.001 | <0.001 |
| ELF1 | 0.0125 | <0.001 | <0.001 |
| FOXA1 | 0.00506 | <0.001 | <0.001 |
| ARID3A | 0.00305 | <0.001 | <0.001 |
| PURA | 0.00299 | <0.001 | <0.001 |
| E2F1 | 0.00299 | <0.001 | <0.001 |
| CEBPZ | 0.00287 | <0.001 | <0.001 |
| YY1 | 0.00267 | <0.001 | <0.001 |
| SPDEF | 0.00251 | <0.001 | <0.001 |
| ZBTB20 | 0.00245 | <0.001 | <0.001 |
| CTCF | 0.00226 | <0.001 | <0.001 |
| FUBP1 | 0.0022 | <0.001 | <0.001 |
| CLOCK | 0.00217 | <0.001 | <0.001 |
| NFIA | 0.00205 | <0.001 | <0.001 |
| SF1 | 0.00202 | <0.001 | <0.001 |
| ZBTB7A | 0.00199 | <0.001 | <0.001 |
| PBX1 | 0.00163 | <0.001 | <0.001 |
| NFIB | 0.00153 | <0.001 | <0.001 |
| CHD1 | 0.00152 | <0.001 | <0.001 |
| FOXM1 | 0.00147 | <0.001 | <0.001 |
| E2F4 | 0.00126 | 0.001 | 0.003 |
| RCOR1 | 0.00122 | <0.001 | <0.001 |
| HINFP | 0.00111 | <0.001 | <0.001 |
| MTF1 | 0.00111 | <0.001 | <0.001 |
| ARNTL | 0.0011 | <0.001 | <0.001 |
| RFX3 | 0.00106 | <0.001 | <0.001 |
| KLF5 | 0.00104 | <0.001 | <0.001 |
| GRHL1 | 0.00103 | 0.002 | 0.004 |
| MYB | 0.00102 | <0.001 | <0.001 |
| GMEB1 | 0.000936 | <0.001 | <0.001 |
| NCALD | 0.000913 | <0.001 | <0.001 |
| E2F2 | 0.000899 | <0.001 | <0.001 |
| NFKB1 | 0.000879 | <0.001 | <0.001 |
| ZSCAN29 | 0.00087 | <0.001 | <0.001 |
| ZBTB43 | 0.000781 | <0.001 | <0.001 |
| FOXJ2 | 0.000744 | 0.003 | 0.006 |
| MSX2 | 0.000639 | <0.001 | 0.002 |
| GATA3 | 0.00058 | <0.001 | <0.001 |
| AHCTF1 | 0.000574 | <0.001 | <0.001 |
| HMG20B | 0.000567 | 0.002 | 0.005 |
| AR | 0.000549 | <0.001 | <0.001 |
| SOX5 | 0.00053 | <0.001 | <0.001 |
| ZFP3 | 0.000523 | <0.001 | <0.001 |
| EBF1 | 0.000507 | <0.001 | <0.001 |
| MYBL2 | 0.000481 | <0.001 | <0.001 |
| RFXANK | 0.000451 | 0.001 | 0.003 |
| NR2E3 | 0.000448 | <0.001 | <0.001 |
| PRRX1 | 0.000446 | <0.001 | <0.001 |
| HOXA10 | 0.000444 | <0.001 | 0.001 |
| FOXP3 | 0.000429 | <0.001 | <0.001 |
| ERF | 0.00042 | 0.003 | 0.007 |
| ALX4 | 0.000374 | <0.001 | <0.001 |
| FOXO3 | 0.000368 | 0.024 | 0.046 |
| ZBTB2 | 0.000366 | 0.011 | 0.023 |
| HOXD10 | 0.000328 | <0.001 | <0.001 |
| HDAC8 | 0.000324 | 0.003 | 0.006 |
| RARB | 0.000299 | <0.001 | <0.001 |
| FOXA3 | 0.000299 | 0.009 | 0.019 |
| MSX1 | 0.000296 | <0.001 | <0.001 |
| FOXJ1 | 0.000265 | <0.001 | 0.002 |
| POU3F1 | 0.000259 | <0.001 | <0.001 |
| HNF4G | 0.000225 | 0.01 | 0.021 |
| SOX10 | 0.000222 | 0.001 | 0.003 |
| SOX18 | 0.000206 | <0.001 | <0.001 |

**B**

| TF | mean diff-reg stat | *p*-val | adj *p*-val |
|---|---:|---:|---:|
| AHDC1 | 0.00463 | <0.001 | <0.001 |
| DRAP1 | 0.00407 | <0.001 | <0.001 |
| BCL6 | 0.00401 | <0.001 | <0.001 |
| CHURC1 | 0.00376 | <0.001 | <0.001 |
| ATF3 | 0.00343 | <0.001 | <0.001 |
| SOX4 | 0.00262 | <0.001 | <0.001 |
| HMGA1 | 0.0025 | <0.001 | <0.001 |
| ETS1 | 0.00208 | <0.001 | <0.001 |
| SP1 | 0.00202 | <0.001 | <0.001 |
| HDAC2 | 0.00201 | <0.001 | <0.001 |
| TAF1 | 0.0019 | <0.001 | <0.001 |
| BHLHE40 | 0.00187 | <0.001 | <0.001 |
| MYC | 0.00176 | <0.001 | <0.001 |
| PHF1 | 0.00173 | <0.001 | <0.001 |
| PDLIM5 | 0.00168 | <0.001 | <0.001 |
| SP2 | 0.00168 | <0.001 | <0.001 |
| MAZ | 0.00164 | <0.001 | <0.001 |
| HOXB2 | 0.00161 | <0.001 | <0.001 |
| ELF3 | 0.00151 | <0.001 | <0.001 |
| ATF7 | 0.0015 | <0.001 | <0.001 |
| BACH1 | 0.00143 | 0.009 | 0.019 |
| ZBTB14 | 0.00111 | 0.002 | 0.006 |
| KLF11 | 0.00111 | <0.001 | <0.001 |
| AHR | 0.00111 | <0.001 | <0.001 |
| MEF2D | 0.0011 | <0.001 | <0.001 |
| PRDM10 | 0.00105 | <0.001 | <0.001 |
| ARNT | 0.00104 | <0.001 | <0.001 |
| FOXC1 | 0.00104 | <0.001 | <0.001 |
| SP100 | 0.00101 | <0.001 | <0.001 |
| HOXC10 | 0.00101 | <0.001 | <0.001 |
| IRF2 | 0.000992 | <0.001 | <0.001 |
| LEF1 | 0.000986 | <0.001 | <0.001 |
| EHF | 0.000973 | 0.012 | 0.023 |
| ZBTB41 | 0.00094 | <0.001 | <0.001 |
| KLF10 | 0.000934 | <0.001 | <0.001 |
| ARID5A | 0.00093 | <0.001 | 0.002 |
| ZSCAN2 | 0.000872 | <0.001 | <0.001 |
| CCDC160 | 0.000839 | 0.002 | 0.004 |
| SOX11 | 0.000834 | <0.001 | <0.001 |
| STAT1 | 0.000768 | 0.013 | 0.025 |
| MEF2A | 0.000744 | <0.001 | <0.001 |
| YOD1 | 0.000733 | <0.001 | <0.001 |
| GMEB2 | 0.000732 | <0.001 | <0.001 |
| HOXB7 | 0.000707 | <0.001 | <0.001 |
| NFE2L1 | 0.000685 | 0.01 | 0.021 |
| RXRA | 0.000664 | <0.001 | <0.001 |
| ATF1 | 0.000662 | 0.002 | 0.006 |
| NR2C2 | 0.000659 | <0.001 | 0.002 |
| ARID3B | 0.00049 | 0.005 | 0.012 |
| CEBPA | 0.000485 | <0.001 | 0.002 |
| ELK3 | 0.000467 | 0.003 | 0.008 |
| TEF | 0.000464 | 0.005 | 0.01 |
| FOSL2 | 0.000461 | <0.001 | 0.001 |
| IKZF3 | 0.00046 | <0.001 | <0.001 |
| HBP1 | 0.000408 | 0.006 | 0.014 |
| RELA | 0.000398 | 0.013 | 0.025 |
| NMRAL1 | 0.000392 | 0.018 | 0.035 |
| EN1 | 0.000372 | <0.001 | <0.001 |
| HES7 | 0.000363 | <0.001 | <0.001 |
| CCNT2 | 0.000361 | 0.012 | 0.024 |
| FOXN3 | 0.000344 | 0.025 | 0.047 |
| ZFP37 | 0.000321 | 0.007 | 0.016 |
| NF1 | 0.000321 | 0.008 | 0.016 |
| BACH2 | 0.000314 | 0.007 | 0.016 |
| SNAPC4 | 0.000314 | 0.004 | 0.01 |
| ABL1 | 0.000275 | 0.003 | 0.007 |
| MNX1 | 0.000259 | <0.001 | 0.002 |
| ZBTB12 | 0.000233 | 0.01 | 0.021 |
| PITX3 | 0.000224 | <0.001 | <0.001 |
| SP5 | 0.000179 | 0.006 | 0.014 |
| ARX | 0.000162 | 0.016 | 0.03 |
| HOXB9 | 0.000158 | <0.001 | <0.001 |
| HOXC13 | 0.000149 | 0.005 | 0.011 |
| HOXA11 | 0.000114 | 0.003 | 0.007 |

***Table S4:*** *Transcription factors with significant differential regulation (FDR $p$-val $<$ 0.05, t-test, Benjamini-Hochberg adjustment) in luminal mature cells that are (a) hypo-regulating and (b) hyper-regulating in BRCA1/2 mutation carriers (BRCAmut) compared to BRCA1/2 wild-type (WT).*

**A**

| TF | mean diff-reg stat | $p$-val | adj $p$-val |
|---|---|---|---|
| EGR1 | 0.0209 | <0.001 | <0.001 |
| BPTF | 0.0122 | <0.001 | <0.001 |
| KLF4 | 0.00629 | <0.001 | <0.001 |
| FOXC1 | 0.0057 | <0.001 | <0.001 |
| ELF1 | 0.00556 | <0.001 | <0.001 |
| ZBTB20 | 0.00526 | <0.001 | <0.001 |
| NFIA | 0.00453 | <0.001 | <0.001 |
| KLF16 | 0.00443 | <0.001 | <0.001 |
| CHD1 | 0.00383 | <0.001 | <0.001 |
| BACH1 | 0.00365 | <0.001 | <0.001 |
| KLF5 | 0.00338 | <0.001 | <0.001 |
| ATF3 | 0.00331 | <0.001 | <0.001 |
| ZBTB14 | 0.0031 | <0.001 | <0.001 |
| SOX11 | 0.00269 | <0.001 | <0.001 |
| PBX1 | 0.00236 | <0.001 | <0.001 |
| BHLHE40 | 0.00235 | <0.001 | <0.001 |
| GLI3 | 0.00219 | <0.001 | <0.001 |
| RCOR1 | 0.00217 | <0.001 | <0.001 |
| EHF | 0.00217 | <0.001 | <0.001 |
| KLF13 | 0.00213 | <0.001 | <0.001 |
| EGR2 | 0.00209 | <0.001 | <0.001 |
| YY1 | 0.00207 | <0.001 | <0.001 |
| ERF | 0.00193 | <0.001 | <0.001 |
| PURA | 0.00186 | <0.001 | <0.001 |
| ELF3 | 0.00179 | <0.001 | <0.001 |
| SOX5 | 0.00176 | <0.001 | <0.001 |
| ZBTB7A | 0.00174 | <0.001 | <0.001 |
| DLX2 | 0.00161 | <0.001 | <0.001 |
| ZBTB11 | 0.00157 | <0.001 | <0.001 |
| AHCTF1 | 0.00157 | <0.001 | <0.001 |
| MAFB | 0.00155 | <0.001 | <0.001 |
| ATF2 | 0.0015 | <0.001 | <0.001 |
| DLX3 | 0.00148 | <0.001 | <0.001 |
| FOXO1 | 0.00146 | <0.001 | <0.001 |
| SOX9 | 0.00145 | <0.001 | <0.001 |
| POU3F1 | 0.00141 | <0.001 | <0.001 |
| HIC1 | 0.0014 | <0.001 | <0.001 |
| E2F1 | 0.00134 | <0.001 | <0.001 |
| ZBTB12 | 0.00132 | <0.001 | <0.001 |
| SPDEF | 0.00131 | <0.001 | <0.001 |
| ALX4 | 0.0013 | <0.001 | <0.001 |
| ELF2 | 0.00129 | <0.001 | <0.001 |
| ALX3 | 0.00111 | <0.001 | <0.001 |
| CEBPA | 0.00108 | <0.001 | <0.001 |
| ID2 | 0.00104 | <0.001 | <0.001 |
| BCL6 | 0.00101 | <0.001 | <0.001 |
| SATB1 | 0.001 | <0.001 | <0.001 |
| HIC2 | 0.000937 | <0.001 | <0.001 |
| RBAK | 0.000929 | <0.001 | <0.001 |
| NCALD | 0.00089 | <0.001 | <0.001 |
| TCFL5 | 0.000862 | <0.001 | <0.001 |
| EGR4 | 0.000854 | <0.001 | <0.001 |
| PRDM10 | 0.000816 | <0.001 | <0.001 |
| SALL2 | 0.000815 | <0.001 | <0.001 |
| EN1 | 0.000741 | <0.001 | <0.001 |
| ASCL1 | 0.000733 | <0.001 | <0.001 |
| RXRA | 0.000732 | <0.001 | <0.001 |
| REST | 0.000727 | <0.001 | <0.001 |
| POU2F1 | 0.000695 | 0.003 | 0.006 |
| KLF2 | 0.000633 | <0.001 | <0.001 |
| MSX2 | 0.000631 | 0.001 | 0.002 |
| KLF10 | 0.00063 | <0.001 | 0.002 |
| FOXL2 | 0.00061 | <0.001 | <0.001 |
| CEBPZ | 0.00061 | 0.024 | 0.035 |
| AHR | 0.000607 | 0.012 | 0.019 |
| IRF4 | 0.000603 | <0.001 | <0.001 |
| ZBTB33 | 0.000597 | 0.012 | 0.018 |
| MYBL1 | 0.00057 | <0.001 | <0.001 |
| PLAG1 | 0.000544 | <0.001 | <0.001 |
| SOX10 | 0.000491 | <0.001 | <0.001 |
| MEF2D | 0.000483 | 0.002 | 0.004 |
| PRDM5 | 0.00047 | 0.003 | 0.005 |
| ZBTB41 | 0.000464 | 0.002 | 0.003 |
| VEZF1 | 0.000446 | 0.016 | 0.024 |
| LCORL | 0.00042 | 0.007 | 0.012 |
| FOXA1 | 0.000413 | <0.001 | <0.001 |
| CRX | 0.000407 | <0.001 | 0.001 |
| EBF1 | 0.000406 | <0.001 | <0.001 |
| ARX | 0.00037 | <0.001 | <0.001 |
| MYB | 0.000355 | <0.001 | <0.001 |
| HLX | 0.000352 | 0.003 | 0.005 |
| HOXC10 | 0.000351 | 0.012 | 0.019 |
| BRCA1 | 0.00035 | 0.004 | 0.007 |
| MNX1 | 0.000327 | <0.001 | <0.001 |
| SIX1 | 0.000314 | <0.001 | <0.001 |
| MEF2C | 0.00031 | 0.009 | 0.015 |
| GLI1 | 0.000302 | <0.001 | <0.001 |
| NKX2-5 | 0.000286 | 0.004 | 0.007 |
| SPI1 | 0.000268 | <0.001 | <0.001 |
| MSX1 | 0.000266 | 0.016 | 0.024 |
| GRHL1 | 0.000261 | 0.023 | 0.034 |
| NR2E3 | 0.000258 | <0.001 | <0.001 |
| HOXB7 | 0.000242 | <0.001 | <0.001 |
| KLF17 | 0.000236 | <0.001 | <0.001 |
| CCDC160 | 0.000229 | <0.001 | <0.001 |
| HOXD9 | 0.000214 | 0.003 | 0.005 |
| EN2 | 0.000196 | 0.003 | 0.005 |
| E2F2 | 0.000189 | <0.001 | <0.001 |
| SIX3 | 0.000178 | 0.006 | 0.009 |
| MYBL2 | 0.000175 | 0.014 | 0.021 |
| SOX6 | 0.000157 | <0.001 | 0.001 |
| SOX18 | 0.000154 | 0.01 | 0.016 |
| OTX1 | 0.000154 | <0.001 | <0.001 |
| ZFP92 | 0.000148 | 0.013 | 0.02 |
| POU3F2 | 0.000147 | 0.003 | 0.005 |
| PAX2 | 0.000128 | <0.001 | <0.001 |
| FOXD3 | 0.000127 | <0.001 | <0.001 |
| DMRT2 | 0.000124 | <0.001 | <0.001 |

**B**

| TF | mean diff-reg stat | $p$-val | adj $p$-val |
|---|---|---|---|
| E2F4 | 0.0171 | <0.001 | <0.001 |
| ELK1 | 0.00997 | <0.001 | <0.001 |
| HDAC2 | 0.00557 | <0.001 | <0.001 |
| MAX | 0.00422 | <0.001 | <0.001 |
| DRAP1 | 0.00385 | <0.001 | <0.001 |
| STAT1 | 0.00368 | <0.001 | <0.001 |
| NFIB | 0.00319 | <0.001 | <0.001 |
| HMGA1 | 0.00292 | <0.001 | <0.001 |
| MAZ | 0.0029 | <0.001 | <0.001 |
| HINFP | 0.00236 | <0.001 | <0.001 |
| CLOCK | 0.00221 | <0.001 | <0.001 |
| DEAF1 | 0.00215 | <0.001 | <0.001 |
| TBP | 0.00211 | <0.001 | <0.001 |
| PARP1 | 0.00204 | <0.001 | <0.001 |
| CNOT3 | 0.00198 | <0.001 | <0.001 |
| HOXA9 | 0.00193 | <0.001 | <0.001 |
| TAF6 | 0.00187 | <0.001 | <0.001 |
| KLF6 | 0.00172 | <0.001 | <0.001 |
| NRF1 | 0.00167 | <0.001 | <0.001 |
| ARNT | 0.00162 | <0.001 | <0.001 |
| ATF1 | 0.0016 | <0.001 | <0.001 |
| ARID5A | 0.00158 | <0.001 | <0.001 |
| ZFP62 | 0.00148 | <0.001 | <0.001 |
| SOX4 | 0.00147 | 0.001 | 0.002 |
| FOXC2 | 0.00133 | <0.001 | <0.001 |
| NFYC | 0.00131 | <0.001 | <0.001 |
| CEBPD | 0.0013 | <0.001 | <0.001 |
| GMEB1 | 0.00121 | <0.001 | <0.001 |
| CXXC1 | 0.00119 | <0.001 | <0.001 |
| IRF2 | 0.00117 | <0.001 | <0.001 |
| TEAD3 | 0.00117 | <0.001 | <0.001 |
| HMG20B | 0.00112 | <0.001 | <0.001 |
| CHURC1 | 0.00109 | <0.001 | <0.001 |
| CTCF | 0.00106 | 0.006 | 0.01 |
| E2F6 | 0.00106 | <0.001 | <0.001 |
| ELK3 | 0.00101 | <0.001 | <0.001 |
| NFIC | 0.001 | <0.001 | <0.001 |
| IRF3 | 0.000984 | <0.001 | <0.001 |
| NR2C2 | 0.000961 | <0.001 | 0.001 |
| STAT3 | 0.000922 | 0.007 | 0.012 |
| PHF1 | 0.000885 | <0.001 | <0.001 |
| MYPOP | 0.000868 | <0.001 | <0.001 |
| HOMEZ | 0.000817 | <0.001 | <0.001 |
| YEATS4 | 0.000798 | <0.001 | <0.001 |
| SREBF2 | 0.00079 | <0.001 | <0.001 |
| TEAD4 | 0.000778 | <0.001 | <0.001 |
| ARNTL | 0.000734 | 0.01 | 0.016 |
| MYC | 0.000731 | 0.002 | 0.003 |
| FUBP1 | 0.000724 | <0.001 | 0.002 |
| DDX20 | 0.000699 | <0.001 | <0.001 |
| PHTF1 | 0.000659 | <0.001 | <0.001 |
| MEF2B | 0.000658 | <0.001 | <0.001 |
| NMRAL1 | 0.000654 | <0.001 | <0.001 |
| GMEB2 | 0.000641 | <0.001 | <0.001 |
| IRF7 | 0.000623 | <0.001 | <0.001 |
| SF1 | 0.000606 | 0.006 | 0.01 |
| MEF2A | 0.000603 | 0.02 | 0.03 |
| CUX1 | 0.000573 | 0.032 | 0.046 |
| MTF1 | 0.000568 | <0.001 | 0.001 |
| TEAD1 | 0.000562 | 0.01 | 0.016 |
| ATF7 | 0.000528 | 0.002 | 0.004 |
| CREB1 | 0.000482 | 0.009 | 0.015 |
| HDAC8 | 0.000442 | <0.001 | <0.001 |
| BRF1 | 0.000439 | 0.022 | 0.033 |
| PRRX2 | 0.000344 | 0.014 | 0.021 |
| HOXC8 | 0.000237 | 0.028 | 0.041 |
| ZSCAN20 | 0.000224 | 0.002 | 0.004 |

***Table S5:*** *Transcription factors with significant differential regulation (FDR* $p\text{-}val < 0.05$*, t-test, Benjamini-Hochberg adjustment) in basal cells that are (a) hypo-regulating and (b) hyper-regulating in BRCA1/2 mutation carriers (BRCAmut) compared to BRCA1/2 wild-type (WT).*

**A**

| TF | mean diff-reg stat | $p$-val | adj $p$-val |
|---|---|---|---|
| CLOCK | 0.0054 | <0.001 | <0.001 |
| STAT1 | 0.00503 | <0.001 | <0.001 |
| BACH1 | 0.00389 | <0.001 | <0.001 |
| HDAC2 | 0.00346 | <0.001 | <0.001 |
| CTCF | 0.00326 | <0.001 | <0.001 |
| KLF13 | 0.00242 | <0.001 | <0.001 |
| ARID3A | 0.0024 | <0.001 | <0.001 |
| IRF4 | 0.00209 | <0.001 | <0.001 |
| BPTF | 0.00209 | <0.001 | <0.001 |
| RBAK | 0.00205 | <0.001 | <0.001 |
| PARP1 | 0.00201 | <0.001 | <0.001 |
| CEBPA | 0.00188 | <0.001 | <0.001 |
| PRRX2 | 0.00181 | <0.001 | <0.001 |
| SP1 | 0.00166 | <0.001 | <0.001 |
| CDX1 | 0.00161 | <0.001 | <0.001 |
| FUBP1 | 0.00159 | <0.001 | <0.001 |
| EGR2 | 0.00157 | <0.001 | <0.001 |
| EHF | 0.00154 | <0.001 | <0.001 |
| FOXM1 | 0.00149 | <0.001 | <0.001 |
| TOPORS | 0.00142 | <0.001 | <0.001 |
| ZBTB7A | 0.00137 | <0.001 | <0.001 |
| GMEB1 | 0.00136 | <0.001 | <0.001 |
| E2F3 | 0.00134 | <0.001 | <0.001 |
| SP100 | 0.00131 | <0.001 | <0.001 |
| GABPA | 0.00131 | <0.001 | <0.001 |
| RCOR1 | 0.00127 | <0.001 | <0.001 |
| ELK4 | 0.00112 | <0.001 | <0.001 |
| NFATC4 | 0.00111 | <0.001 | <0.001 |
| STAT4 | 0.00111 | <0.001 | <0.001 |
| FOXJ2 | 0.00109 | <0.001 | <0.001 |
| ERF | 0.00103 | <0.001 | <0.001 |
| TAF1 | 0.00102 | <0.001 | <0.001 |
| TCFL5 | 0.000987 | <0.001 | <0.001 |
| NCALD | 0.000986 | <0.001 | <0.001 |
| NFE2L1 | 0.000889 | <0.001 | <0.001 |
| ELK3 | 0.000882 | <0.001 | <0.001 |
| E2F2 | 0.000857 | <0.001 | <0.001 |
| CEBPZ | 0.000835 | <0.001 | <0.001 |
| RFX2 | 0.000833 | <0.001 | <0.001 |
| VEZF1 | 8e-04 | <0.001 | <0.001 |
| DLX3 | 0.000798 | <0.001 | <0.001 |
| NR2C2 | 0.000796 | <0.001 | <0.001 |
| NFATC3 | 0.000788 | <0.001 | <0.001 |
| ZSCAN29 | 0.000778 | <0.001 | <0.001 |
| HIC2 | 0.000771 | <0.001 | <0.001 |
| FOXL2 | 0.000769 | <0.001 | <0.001 |
| CRX | 0.00076 | <0.001 | <0.001 |
| AHCTF1 | 0.000741 | <0.001 | <0.001 |
| ETS1 | 0.000724 | 0.012 | 0.022 |
| NR4A2 | 0.000719 | <0.001 | <0.001 |
| BACH2 | 0.000719 | <0.001 | 0.001 |
| POU2F1 | 0.000711 | <0.001 | <0.001 |
| SATB1 | 0.00071 | <0.001 | <0.001 |
| ZBTB37 | 0.000688 | <0.001 | <0.001 |
| CNOT3 | 0.000677 | <0.001 | 0.002 |
| BRCA1 | 0.000658 | <0.001 | <0.001 |
| TBP | 0.000657 | 0.002 | 0.004 |
| ZKSCAN8 | 0.00065 | <0.001 | <0.001 |
| ARID5A | 0.000649 | <0.001 | <0.001 |
| ALX3 | 6e-04 | 0.001 | 0.003 |
| E2F8 | 0.000591 | <0.001 | <0.001 |
| KLF16 | 0.000589 | <0.001 | <0.001 |
| GLI3 | 0.000535 | 0.002 | 0.005 |
| ZBTB11 | 0.000533 | 0.004 | 0.009 |
| ZEB1 | 0.000531 | 0.006 | 0.012 |
| ARID3B | 0.000519 | <0.001 | <0.001 |
| SIX3 | 5e-04 | <0.001 | <0.001 |
| GMEB2 | 0.000489 | 0.001 | 0.003 |
| ASCL1 | 0.000474 | <0.001 | <0.001 |
| ATF7 | 0.000459 | 0.003 | 0.007 |
| SPDEF | 0.000425 | <0.001 | <0.001 |
| KLF2 | 0.000425 | 0.006 | 0.012 |
| ZFP92 | 0.000396 | <0.001 | <0.001 |
| FOXC2 | 0.000377 | 0.009 | 0.017 |
| OTX1 | 0.000371 | <0.001 | <0.001 |
| ZFP3 | 0.000369 | <0.001 | <0.001 |
| HOXA13 | 0.000339 | <0.001 | 0.002 |
| IKZF1 | 0.000332 | <0.001 | <0.001 |
| NR2E3 | 0.00033 | <0.001 | <0.001 |
| PURG | 0.000286 | 0.006 | 0.011 |
| FOXP3 | 0.00028 | 0.005 | 0.01 |
| BCL6B | 0.000278 | <0.001 | <0.001 |
| IKZF3 | 0.000255 | 0.009 | 0.016 |
| IRF6 | 0.000214 | 0.005 | 0.01 |
| GATA3 | 0.000193 | 0.01 | 0.018 |
| PATZ1 | 0.000191 | 0.02 | 0.035 |
| MYBL2 | 0.000181 | <0.001 | <0.001 |
| MYB | 0.000145 | 0.029 | 0.05 |
| SOX11 | 0.000139 | 0.012 | 0.021 |
| CCDC160 | 0.000139 | 0.01 | 0.018 |
| POU4F3 | 0.000127 | <0.001 | <0.001 |
| PAX2 | 0.000126 | <0.001 | <0.001 |
| NKX2-5 | 9.87e-05 | 0.003 | 0.007 |
| ARX | 8.32e-05 | <0.001 | <0.001 |

**B**

| TF | mean diff-reg stat | $p$-val | adj $p$-val |
|---|---|---|---|
| EGR1 | 0.0122 | <0.001 | <0.001 |
| CHURC1 | 0.00335 | <0.001 | <0.001 |
| ELF1 | 0.00319 | <0.001 | <0.001 |
| E2F4 | 0.00284 | <0.001 | <0.001 |
| ELK1 | 0.00261 | <0.001 | <0.001 |
| ARNTL | 0.00245 | <0.001 | <0.001 |
| ATF2 | 0.0021 | <0.001 | <0.001 |
| HMGA1 | 0.00192 | <0.001 | <0.001 |
| CEBPB | 0.00183 | <0.001 | <0.001 |
| DEAF1 | 0.00176 | <0.001 | <0.001 |
| BCLAF1 | 0.00176 | <0.001 | 0.002 |
| BCL6 | 0.00171 | <0.001 | <0.001 |
| KLF5 | 0.00159 | <0.001 | <0.001 |
| PDLIM5 | 0.00143 | <0.001 | <0.001 |
| ZBTB14 | 0.0014 | <0.001 | <0.001 |
| ARNT | 0.00138 | <0.001 | <0.001 |
| MAX | 0.00133 | <0.001 | <0.001 |
| DRAP1 | 0.00129 | <0.001 | <0.001 |
| STAT5A | 0.00115 | <0.001 | <0.001 |
| IRF7 | 0.00114 | <0.001 | <0.001 |
| MAZ | 0.0011 | 0.001 | 0.002 |
| STAT3 | 0.0011 | <0.001 | <0.001 |
| KLF11 | 0.000963 | <0.001 | <0.001 |
| ZBTB2 | 0.000956 | <0.001 | <0.001 |
| TWIST1 | 0.000894 | <0.001 | <0.001 |
| KLF9 | 0.000885 | <0.001 | <0.001 |
| MEF2B | 0.000854 | <0.001 | <0.001 |
| LEF1 | 0.000847 | <0.001 | <0.001 |
| HSF1 | 0.00078 | <0.001 | <0.001 |
| RAB18 | 0.000747 | <0.001 | <0.001 |
| KLF10 | 0.000722 | 0.002 | 0.004 |
| HOXD10 | 0.000708 | <0.001 | <0.001 |
| PRRX1 | 0.000683 | <0.001 | 0.001 |
| SF1 | 0.000668 | <0.001 | <0.001 |
| RARB | 0.000657 | <0.001 | <0.001 |
| SREBF2 | 0.000654 | <0.001 | <0.001 |
| SOX4 | 0.000628 | <0.001 | <0.001 |
| ILF2 | 0.000622 | <0.001 | <0.001 |
| ID2 | 0.000606 | 0.002 | 0.005 |
| MAFB | 0.000538 | <0.001 | <0.001 |
| TEAD4 | 0.000531 | <0.001 | <0.001 |
| YY1 | 0.000525 | 0.004 | 0.009 |
| IRF2 | 0.000508 | 0.023 | 0.039 |
| MSX2 | 0.000489 | <0.001 | <0.001 |
| YOD1 | 0.00048 | <0.001 | <0.001 |
| CXXC1 | 0.000477 | 0.007 | 0.013 |
| KLF15 | 0.000455 | <0.001 | <0.001 |
| POU3F1 | 0.000441 | <0.001 | <0.001 |
| MSX1 | 0.000408 | 0.016 | 0.029 |
| MEF2D | 0.00039 | 0.005 | 0.01 |
| IRF1 | 0.000379 | 0.028 | 0.049 |
| DMRT2 | 0.000369 | 0.008 | 0.015 |
| HOXC8 | 0.000346 | 0.013 | 0.023 |
| ZBTB26 | 0.00033 | 0.002 | 0.004 |
| GRHL1 | 0.000327 | 0.004 | 0.008 |
| HOXB2 | 0.000284 | 0.01 | 0.018 |
| MEF2C | 0.000284 | 0.017 | 0.03 |
| FOXO6 | 0.000253 | 0.003 | 0.006 |
| HOXD3 | 0.000245 | 0.006 | 0.012 |
| HOXA3 | 0.000228 | 0.006 | 0.012 |
| EN2 | 0.000213 | 0.02 | 0.035 |
| MECOM | 0.000209 | 0.005 | 0.01 |
| POU3F2 | 2e-04 | <0.001 | 0.002 |
| SOX18 | 0.000181 | 0.012 | 0.022 |
| E2F7 | 0.000164 | 0.002 | 0.004 |
| FOXD3 | 0.000164 | <0.001 | <0.001 |
| CEBPE | 0.000161 | 0.002 | 0.005 |
| ISL1 | 0.000153 | <0.001 | 0.002 |

***Table S6:*** *Transcription factors with significant differential regulation (FDR $p\text{-}val < 0.05$, t-test, Benjamini-Hochberg adjustment) in fibroblasts (type 1) that are (a) hypo-regulating and (b) hyper-regulating in BRCA1/2 mutation carriers (BRCAmut) compared to BRCA1/2 wild-type (WT).*

**A**

| TF | mean diff-reg stat | *p*-val | adj *p*-val |
|---|---|---|---|
| BCLAF1 | 0.00631 | <0.001 | <0.001 |
| BPTF | 0.00453 | <0.001 | <0.001 |
| ELF1 | 0.00374 | <0.001 | <0.001 |
| KLF4 | 0.00355 | <0.001 | <0.001 |
| CLOCK | 0.00275 | <0.001 | <0.001 |
| PURA | 0.00263 | <0.001 | <0.001 |
| CTCF | 0.00233 | <0.001 | <0.001 |
| FOXC1 | 0.00204 | <0.001 | <0.001 |
| CEBPZ | 0.0019 | <0.001 | <0.001 |
| ZBTB14 | 0.00189 | <0.001 | <0.001 |
| EHF | 0.00182 | <0.001 | <0.001 |
| STAT1 | 0.00166 | <0.001 | <0.001 |
| ARID3A | 0.00165 | 0.002 | 0.005 |
| CEBPA | 0.00157 | <0.001 | <0.001 |
| DLX2 | 0.00155 | <0.001 | <0.001 |
| PARP1 | 0.00142 | <0.001 | <0.001 |
| E2F1 | 0.00134 | <0.001 | <0.001 |
| PRRX2 | 0.00132 | <0.001 | <0.001 |
| MAZ | 0.00131 | <0.001 | 0.001 |
| ATF3 | 0.00122 | 0.005 | 0.011 |
| DLX3 | 0.0012 | <0.001 | <0.001 |
| ZBTB20 | 0.0012 | <0.001 | <0.001 |
| CUX1 | 0.0012 | <0.001 | <0.001 |
| SF1 | 0.00114 | <0.001 | <0.001 |
| NFATC1 | 0.00112 | <0.001 | <0.001 |
| CDX1 | 0.0011 | <0.001 | <0.001 |
| EGR2 | 0.00108 | <0.001 | <0.001 |
| ELF3 | 0.00105 | <0.001 | <0.001 |
| KLF2 | 0.000999 | <0.001 | <0.001 |
| IRF3 | 0.000943 | <0.001 | <0.001 |
| KLF13 | 0.000922 | 0.001 | 0.003 |
| CREB1 | 0.000909 | <0.001 | <0.001 |
| EGR4 | 0.000906 | <0.001 | <0.001 |
| CHD1 | 0.000896 | <0.001 | <0.001 |
| TOPORS | 0.000781 | <0.001 | <0.001 |
| NR4A2 | 0.000743 | <0.001 | <0.001 |
| ELK3 | 0.000715 | 0.002 | 0.005 |
| CDC5L | 0.00071 | 0.007 | 0.015 |
| YY1 | 0.000684 | 0.002 | 0.005 |
| HOXC8 | 0.000642 | <0.001 | <0.001 |
| STAT4 | 0.000624 | 0.001 | 0.004 |
| ABL1 | 0.000597 | <0.001 | 0.002 |
| ALX4 | 0.000593 | <0.001 | <0.001 |
| ZSCAN29 | 0.000561 | <0.001 | <0.001 |
| ZBTB11 | 0.000559 | 0.001 | 0.003 |
| KLF6 | 0.000545 | 0.005 | 0.012 |
| SIX3 | 0.000538 | <0.001 | <0.001 |
| OVOL1 | 0.00051 | <0.001 | <0.001 |
| SOX18 | 0.000489 | <0.001 | <0.001 |
| E2F2 | 0.000466 | <0.001 | <0.001 |
| DLX4 | 0.000461 | <0.001 | <0.001 |
| HOXA13 | 0.000455 | <0.001 | <0.001 |
| ZBTB7A | 0.000455 | 0.006 | 0.014 |
| ZBTB2 | 0.000443 | 0.001 | 0.003 |
| VEZF1 | 0.000443 | 0.019 | 0.037 |
| ID2 | 0.000426 | 0.017 | 0.034 |
| HMG20A | 0.000419 | 0.002 | 0.005 |
| ZIC1 | 0.000418 | <0.001 | <0.001 |
| ZBTB12 | 0.000416 | <0.001 | <0.001 |
| HOXA10 | 0.000405 | 0.008 | 0.016 |
| FOXC2 | 0.000394 | 0.002 | 0.004 |
| NFATC3 | 0.000389 | <0.001 | <0.001 |
| DBX2 | 0.000378 | <0.001 | <0.001 |
| NCALD | 0.000376 | <0.001 | <0.001 |
| FOXJ3 | 0.000364 | 0.006 | 0.013 |
| HOXA11 | 0.000337 | <0.001 | <0.001 |
| SIX5 | 0.000337 | 0.006 | 0.014 |
| HES7 | 0.000318 | 0.012 | 0.024 |
| BRCA1 | 0.000302 | 0.003 | 0.007 |
| IKZF1 | 0.000302 | <0.001 | <0.001 |
| IRF6 | 0.00029 | 0.001 | 0.003 |
| FOXA1 | 0.000289 | 0.019 | 0.037 |
| SOX11 | 0.000276 | <0.001 | <0.001 |
| OVOL2 | 0.000246 | <0.001 | <0.001 |
| BCL6B | 0.000242 | <0.001 | 0.003 |
| PURG | 0.000232 | 0.002 | 0.006 |
| DMRT2 | 0.000182 | 0.015 | 0.03 |
| BARX2 | 0.000141 | 0.024 | 0.047 |
| NKX2-5 | 0.000122 | <0.001 | <0.001 |
| MECOM | 0.000101 | 0.024 | 0.046 |
| KLF17 | 9.16e-05 | 0.015 | 0.03 |
| PAX2 | 8.86e-05 | 0.023 | 0.045 |
| HAND1 | 6.32e-05 | 0.025 | 0.047 |
| OTX1 | 5.86e-05 | 0.004 | 0.009 |

**B**

| TF | mean diff-reg stat | *p*-val | adj *p*-val |
|---|---|---|---|
| ELK1 | 0.00742 | <0.001 | <0.001 |
| E2F4 | 0.0053 | <0.001 | <0.001 |
| MAX | 0.00325 | <0.001 | <0.001 |
| KLF9 | 0.00277 | <0.001 | <0.001 |
| ARNT | 0.00276 | <0.001 | <0.001 |
| DRAP1 | 0.00272 | <0.001 | <0.001 |
| BCL6 | 0.00261 | <0.001 | <0.001 |
| CEBPB | 0.0024 | <0.001 | <0.001 |
| CHURC1 | 0.00194 | <0.001 | <0.001 |
| MEF2D | 0.00179 | <0.001 | <0.001 |
| ZFHX3 | 0.00175 | <0.001 | <0.001 |
| PDLIM5 | 0.00167 | <0.001 | <0.001 |
| MEF2A | 0.00161 | <0.001 | <0.001 |
| ARNTL | 0.00151 | <0.001 | <0.001 |
| BACH1 | 0.00145 | 0.006 | 0.014 |
| SP1 | 0.00143 | <0.001 | <0.001 |
| SREBF2 | 0.00141 | <0.001 | <0.001 |
| HDAC2 | 0.00122 | <0.001 | <0.001 |
| MSC | 0.00121 | <0.001 | <0.001 |
| ETS1 | 0.00119 | <0.001 | <0.001 |
| NRF1 | 0.00111 | <0.001 | <0.001 |
| AHR | 0.00107 | <0.001 | <0.001 |
| HSF1 | 0.00105 | <0.001 | <0.001 |
| ILF2 | 0.000993 | <0.001 | <0.001 |
| TBP | 0.000991 | <0.001 | 0.001 |
| ATF2 | 0.000987 | <0.001 | <0.001 |
| IRF9 | 0.000986 | <0.001 | <0.001 |
| MECP2 | 0.000945 | <0.001 | <0.001 |
| STAT5A | 0.000907 | <0.001 | <0.001 |
| LEF1 | 0.00088 | <0.001 | <0.001 |
| KLF5 | 0.000843 | <0.001 | 0.002 |
| CNOT3 | 0.000838 | <0.001 | <0.001 |
| TAF1 | 0.000833 | <0.001 | <0.001 |
| SP2 | 0.000822 | 0.002 | 0.005 |
| KLF11 | 0.000804 | <0.001 | 0.001 |
| ZBTB41 | 0.000737 | <0.001 | <0.001 |
| YOD1 | 0.000715 | <0.001 | <0.001 |
| MEF2B | 7e-04 | <0.001 | <0.001 |
| NFATC4 | 0.000682 | <0.001 | <0.001 |
| NR2C2 | 0.000676 | <0.001 | 0.001 |
| E2F6 | 0.000664 | <0.001 | <0.001 |
| GMEB2 | 0.000636 | <0.001 | <0.001 |
| STAT3 | 0.000636 | 0.001 | 0.004 |
| RAB18 | 0.00063 | <0.001 | <0.001 |
| PBX1 | 0.000626 | <0.001 | 0.002 |
| BACH2 | 0.000604 | 0.002 | 0.005 |
| KLF15 | 0.000595 | <0.001 | <0.001 |
| EN2 | 0.000592 | <0.001 | <0.001 |
| ATF7 | 0.000563 | <0.001 | 0.001 |
| POU2F1 | 0.000543 | 0.009 | 0.019 |
| TFE3 | 0.00047 | 0.002 | 0.005 |
| NRL | 0.000441 | <0.001 | <0.001 |
| ELK4 | 0.000416 | 0.013 | 0.026 |
| TWIST1 | 0.000396 | 0.017 | 0.035 |
| SNAPC5 | 0.000373 | 0.003 | 0.007 |
| ASCL1 | 0.00037 | <0.001 | <0.001 |
| HLX | 0.000361 | 0.012 | 0.024 |
| POU3F1 | 0.000319 | 0.007 | 0.016 |
| HOXD10 | 0.000306 | 0.006 | 0.014 |
| ZFP92 | 0.000257 | 0.022 | 0.044 |
| MYB | 0.00025 | 0.007 | 0.016 |

***Table S7:*** *Transcription factors with significant differential regulation (FDR $p\text{-}val < 0.05$, t-test, Benjamini-Hochberg adjustment) in fibroblasts (type 2) that are (a) hypo-regulating and (b) hyper-regulating in BRCA1/2 mutation carriers (BRCAmut) compared to BRCA1/2 wild-type (WT).*

| | *HR* (95%CI) | *p*-value |
|---|---|---|
| cg10020083 methylation | 0.764 (0.395 - 1.48) | 0.423 |
| Age (years) | 1.53 (1.04 - 2.23) | 0.029 |
| AJCC Stage (III-IV vs. I-II) | 2.99 (1.37 - 6.52) | 0.006 |
| Fibroblast fraction | 0.37 (0.006 - 24.2) | 0.641 |
| Fat cell fraction | 3.2 (0.1 - 102) | 0.511 |
| Immune cell fraction | 1.1 (0.061 - 19.7) | 0.948 |

| | *HR* (95%CI) | *p*-value |
|---|---|---|
| cg23403350 methylation | 1.59 (1.2 - 2.12) | 0.001 |
| Age (years) | 1.65 (1.12 - 2.41) | 0.01 |
| AJCC Stage (III-IV vs. I-II) | 3.83 (1.72 - 8.54) | 0.001 |
| Fibroblast fraction | 0.239 (0.004 - 15.6) | 0.502 |
| Fat cell fraction | 1.19 (0.037 - 38.7) | 0.921 |
| Immune cell fraction | 0.744 (0.04 - 13.8) | 0.842 |

| | *HR* (95%CI) | *p*-value |
|---|---|---|
| cg04110559 methylation | 1.67 (1.07 - 2.6) | 0.023 |
| Age (years) | 1.64 (1.12 - 2.41) | 0.011 |
| AJCC Stage (III-IV vs. I-II) | 4.02 (1.75 - 9.24) | 0.001 |
| Fibroblast fraction | 0.206 (0.003 - 12.2) | 0.448 |
| Fat cell fraction | 1.69 (0.06 - 47) | 0.758 |
| Immune cell fraction | 0.814 (0.043 - 15.4) | 0.891 |

***Table S8:*** *Survival analyses (Cox regression) showing association of CpG methylation level with patient survival outcome in the TCGA BRCA dataset [61] for the 3 CpGs annotated to the PRR7 promoter (Figure 8) on the Illumina 450K DNAme microarray, adjusted for confounding by cell-type heterogeneity and clinical covariates.*

| | | | |
|---|---|---|---|
| E6.10.1046 | E6.17.1588 | E6.9.1024 | E7.17.1348 |
| E6.10.1048 | E6.17.1611 | E7.10.760 | E7.17.1353 |
| E6.10.1049 | E6.17.1612 | E7.10.761 | E7.6.262 |
| E6.10.1050 | E6.17.1617 | E7.10.762 | E7.6.264 |
| E6.10.1051 | E6.22.1852 | E7.10.764 | E7.8.311 |
| E6.10.1052 | E6.22.1866 | E7.10.766 | E7.8.312 |
| E6.10.1055 | E6.8.791 | E7.10.768 | E7.8.317 |
| E6.10.1056 | E6.8.792 | E7.12.858 | E7.8.318 |
| E6.10.1057 | E6.8.794 | E7.12.861 | E7.8.329 |
| E6.10.1058 | E6.8.801 | E7.12.869 | E7.8.333 |
| E6.12.1274 | E6.8.802 | E7.12.871 | E7.8.343 |
| E6.12.1289 | E6.8.813 | E7.14.895 | E7.9.547 |
| E6.13.1378 | E6.8.815 | E7.17.1334 | E7.9.554 |
| E6.13.1381 | E6.8.816 | E7.17.1335 | E7.9.556 |
| E6.13.1389 | E6.8.820 | E7.17.1342 | E7.9.562 |
| E6.16.1501 | E6.8.821 | E7.17.1346 | E7.9.569 |
| E6.17.1586 | E6.9.1022 | E7.17.1347 | E7.9.571 |

***Table S9:*** *The 68 epiblast cells identified previously [84] and used in this work, identified as cluster . The listed cell IDs correspond to the IDs given in the original study that generated these data [42].*

| | | | |
|---|---|---|---|
| E5.12.1038 | E5.14.1812 | E5.37.3243 | E6.10.1043 |
| E6.10.1044 | E6.10.1045 | E6.10.1060 | E6.10.1061 |
| E6.12.1273 | E6.12.1279 | E6.12.1281 | E6.13.1382 |
| E6.18.1641 | E7.10.767 | E7.11.841 | E7.11.842 |
| E7.11.843 | E7.11.845 | E7.11.847 | E7.11.855 |
| E7.12.857 | E7.12.866 | E7.12.872 | E7.13.882 |
| E7.E7.13.883 | E7.13.884 | E7.13.892 | E7.14.894 |
| E7.14.904 | E7.14.905 | E7.14.906 | E7.14.908 |
| E7.16.1135 | E7.16.1169 | E7.8.327 | E7.9.539 |
| E7.9.550 | | | |

***Table S10:*** *The 37 PE (primitive endoderm) cells identified as cluster 4 in Figure S2. The listed cell IDs correspond to the IDs given in the original study that generated these data [42].*
